# Cross-Comparison of Gut Metagenomic Profiling Strategies

**DOI:** 10.1101/2023.11.25.568646

**Authors:** Gábor Gulyás, Balázs Kakuk, Ákos Dörmő, Tamás Járay, István Prazsák, Zsolt Csabai, Miksa Máté Henkrich, Zsolt Boldogkői, Dóra Tombácz

**Author notes:** The first three authors contributed equally to this work. Correspondence &.

## Abstract

A critical issue in microbiome research is the selection of reliable laboratory and bioinformatics pipelines. In the absence of generally accepted technical benchmarks and evaluation standards, comparing data generated by different studies becomes challenging. In this work, we carried out the most comprehensive study to date on this topic. We encompassed every stage of processing, from DNA extraction to computational assessment. We adopted four procedures for DNA purification, six for library construction, three for sequencing, and five for bioinformatics. Additionally, we used datasets published by others to corroborate our results. We introduced a software tool that distinctively delivers consistent results, irrespective of sample or dataset origins. This study underscores the importance of methodological optimization at the outset of research projects to ensure the reliability of results and their comparability with findings from other studies. Additionally, this study provides an optimized robust pipeline for gut microbiome analysis.

## INTRODUCTION

Metagenomics is an emerging discipline that has recently experienced a boom due to rapid advancements in DNA and RNA sequencing, as well as associated technologies in molecular biology, genomics, and bioinformatics^1,2^. Current investigations using innovative approaches have not only unveiled the vast diversity and complexity of the gut microbiome but also underscored its critical role in human health and disease, enhancing our comprehension of the dynamic interactions within microbial communities and between the hosts and their microbes^3,5,6,7,8^. Cutting-edge sequencing technologies now enable comprehensive taxonomic and functional profiling of microbial communities. Recent nucleic acid purification methods are efficient, library preparation methods yield high-quality samples, and current software tools adeptly handle metagenomics data. Although a plethora of metagenomics methods are available for both wet and dry lab steps, there have been limited standardization initiatives to date^9,10^. While there are excellent works that address the impact of sequencing techniques and bioinformatics strategies on the variability of results^2,11,12,13^ remarkably, no comprehensive study simultaneously assesses both the complete laboratory processes and the subsequent data analysis protocols (**Supplementary Data 1**).

Choosing the best DNA purification method^14,15,16,17,18,19^, sequencing platform, shotgun vs. amplicon sequencing^20,21^, optimal 16S rRNA hypervariable region^22,23^, and appropriate data analysis software/database is crucial (**Supplementary Data 1**). An inappropriate choice of DNA extraction kit might result in inefficient lysis of the cell walls of Gram-positive bacteria, leading to underrepresentation of species with more rigid cell wall structures^24,25,26,27^. Indeed, technical variability among studies is frequently attributed to differences in DNA isolation methods^28,29^. Similarly, the selection of library preparation protocols has been demonstrated to be vital, given the notable differences in taxonomic accuracy among these methods^12,30,31^.

The impact of sequencing techniques^2^ and bioinformatics strategies on the variability of results is seldom addressed in scientific literature. However, choosing the right bioinformatics techniques and databases is essential, as they influence the results of community composition^32,33^. While there is a wide range of bioinformatics methods, no single workflow is ideal for all types of sequencing data.

Two primary sequencing methods are short-read sequencing (SRS) and long-read sequencing (LRS)^34^. Traditionally, metagenomic whole genome sequencing (mWGS) is conducted on SRS platforms, accompanied by popular bioinformatic tools like Kraken2^35,36,37^, sourmash^38,39^, and MEGAN^40,41^. A recent benchmark, using sequencing datasets of different mock microbial communities, showed that sourmash is the only tool that was able to produce excellent accuracy and precision on both SRS and LRS data^33^.

In another study, where the authors compared different programs and databases for 16S-Seq^42^, a conclusion was that Kraken2, while originally developed for the analysis of WGS reads, proved to be applicable to a non-16S database (RefSeq).

The emergence of LRS platforms from Oxford Nanopore Technologies (ONT) and Pacific Biosciences (PacBio) has allowed the sequencing of the full 16S, potentially enabling higher resolution of classification^12^. However, there are limited bioinformatic tools for this context, with Emu and EPI2ME^11^ being significant exceptions.

Recent findings show a strong similarity between the canine and the human gut microbiomes^43^. Given the genetic uniformity within dog breeds and the ability to control their diet, dogs stand out as an ideal model for microbiome research, with findings potentially applicable to human contexts.

In our work, we examined the fecal samples from a Hungarian breed known as Pumi. Aiming to comprehensively evaluate metagenomic workflows for gut microbiome profiling, we performed a series of tests comparing several techniques, including DNA isolation, library preparation, sequencing, and bioinformatics approaches in fecal microbial analysis. We created a new versatile program, named minitax, designed to reduce variability in bioinformatics workflows and provide uniform analysis across various sequencing platforms.

In summary, the main objectives of our work were to evaluate existing wet-lab and dry-lab techniques, identify the best practices for each stage of the process to ensure reliable and consistent gut microbiome profiling, and to develop a universally applicable bioinformatic tool.

## RESULTS

### 1. Study Design

The objective of this study was to evaluate how various protocols influence the results concerning the microbial composition of the gut microbiome. These protocols cover DNA isolation, library preparation, sequencing, and bioinformatics techniques (**Fig. 1**). We evaluated the quantity, quality, and reproducibility of DNA extracted by various isolation kits (**Supplementary Data 2a**). We also characterized the microbial composition, examining predominant taxa and overall diversity. Libraries were prepared for Illumina mWGS using the DNA Prep kit and for amplicon sequencing of V-regions V1-V3, V1-V2, and V3-V4 of the 16S rRNA gene (**Supplementary Data 2b**). We also prepared V1-V9 libraries spanning the entire 16S rRNA region for sequencing on two LRS platforms: ONT MinION and PacBio Sequel IIe. Libraries were assessed for quality, volume, and consistency.

**Fig. 1:**
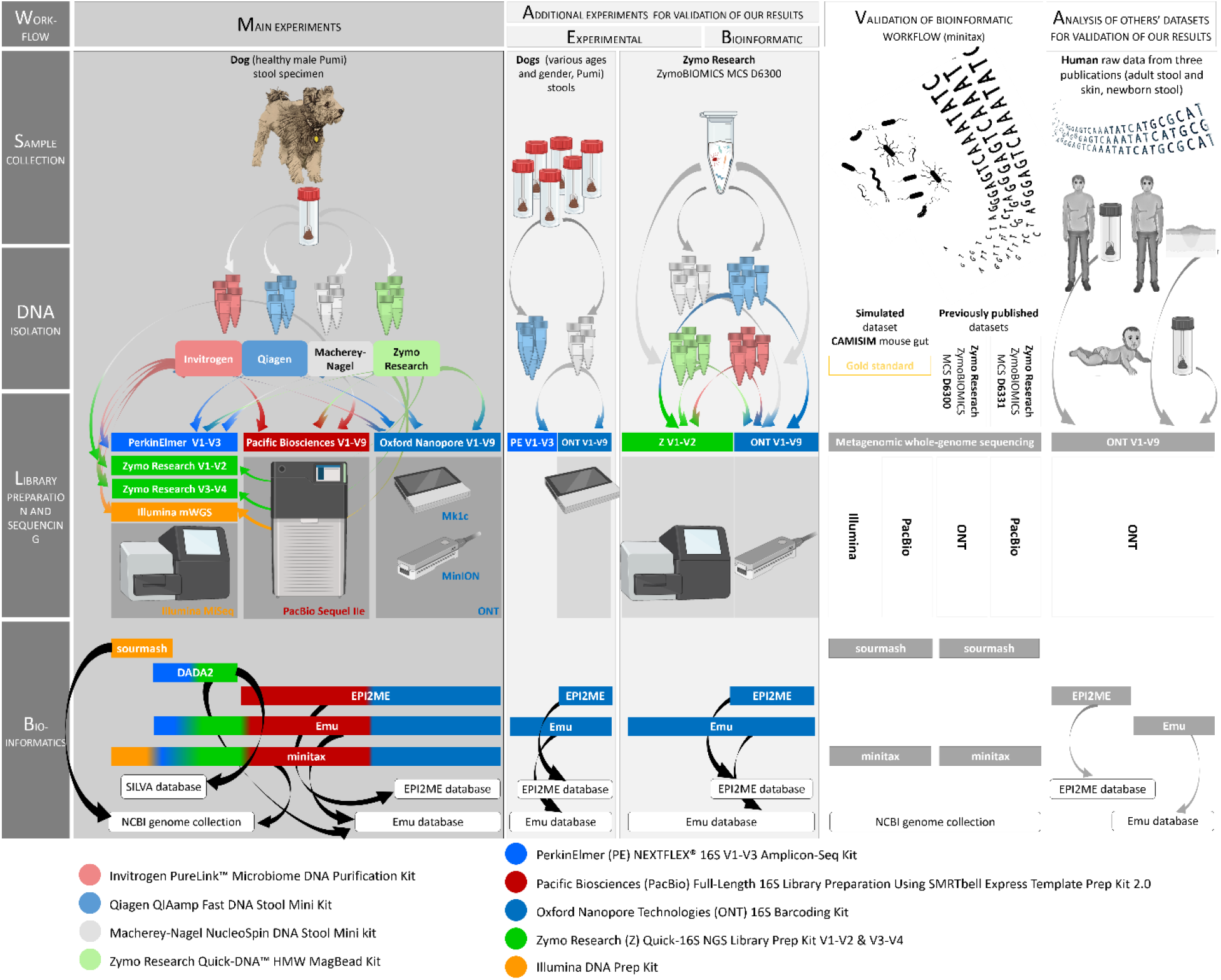
Workflow of the study. The figure provides a detailed representation of the workflows conducted in this study. The figure illustrates the workflows conducted in this study, which included: 1) evaluating the efficacy of various DNA isolation kits in terms of quality, quantity, and microbial representation from canine stool samples; 2) comparing library preparation techniques on SRS and LRS platforms for reproducibility; 3) introducing “minitax”, a tool designed to ensure consistent analysis across multiple sequencing platforms; 4) assessing the influence of different databases and tools on microbial profiling; and 5) comparing 16S V1-V9 sequencing on ONT and PacBio platforms to address literature gaps and emphasize bioinformatics workflows. Our goal was to identify reliable procedures for robust and reproducible gut microbiome profiling across both wet-lab and dry-lab methodologies. We performed additional experiments to validate the most extreme experimental and bioinformatics results, particularly focusing on methods that yielded the most inconsistent outcomes in comparison to other techniques. For this purpose, we utilized samples from six additional dogs of various genders and ages. The workflow involved: 1) DNA isolation using the Qiagen kit, 2) Library preparation with the PerkinElmer V1-V3 kit, and 3) Analysis of V1-V9 libraries using the EPI2ME software. Furthermore, we carried out experiments employing a Microbial Community Standard (MCS; Zymo Research D6300) to validate the effectiveness of the four DNA isolation kits used. This included: 1) DNA isolation using the kits applied for the dog samples, 2) Preparation of V1-V2 and V1-V9 libraries, and 3) Sequencing on the corresponding Illumina and ONT platforms based on the library. Additionally, we conducted an *in silico* analysis to validate our bioinformatic tool, minitax. We compared the performance of minitax and sourmash by utilizing them for the analysis of: 1) simulated PacBio and Illumina data^44^, and 2) previously published datasets^33^. For further validation of our results, we also used previously published metagenomics data from human sources encompassing skin^57^ as well as fecal samples from newborns^55^ and adults^56^.

We employed the following bioinformatics approaches: DADA2^45^ for amplicon-based SRS datasets, sourmash^46^ for WGS samples, and Emu, a recently published, highly accurate software, optimized for LRS 16S-Seq^47^. We also utilized the ONT’s company-specific pipeline, EPI2ME^48^ for processing the nanopore data.

To assess how library preparation methods, affect microbial composition accuracy, we used the same DNA isolation kit to standardize samples and eliminate potential inconsistencies. We devised a versatile, universally applicable program, called ‘minitax’, based on the minimap2 aligner^49^ to process various datasets, ensuring precise taxonomic assignments for both sequencing types and platforms (**Supplementary Data 2c**). To compare and validate the results, different stages of the process were evaluated using a variety of samples and data types (**Fig. 1**).

### 2. Comparison of DNA Preparation Techniques

Most DNA isolation kits employ affinity-based DNA purification, inhibitor removal buffers or columns, and lysis buffers, enzymes, or bead-beating for cell wall disruption. Although several studies advocate bead-beating for this purpose^24,26^, many popular commercial kits do not include this step. We assessed four commercial DNA isolation kits differing in the aforementioned techniques (**Supplementary Data 2a**). Among the four methodologies scrutinized, the Zymo approach necessitated the most extensive hands-on time, distinguishing it from the others in terms of labor intensity (**Supplementary Fig. 1**). The remaining three methods exhibited parity in this respect. The DNA extraction kits used in this study showed significant differences in both the quantity and quality of the extracted nucleic acid (**Supplementary Data 3**), as evidenced by variations in yield (**Figs. 2a-b**, **Supplementary Fig. 2**), microbe-to-host ratio (**Figs. 2c-d**), and reproducibility (**Figs. 2e-f**).

**Fig. 2:**
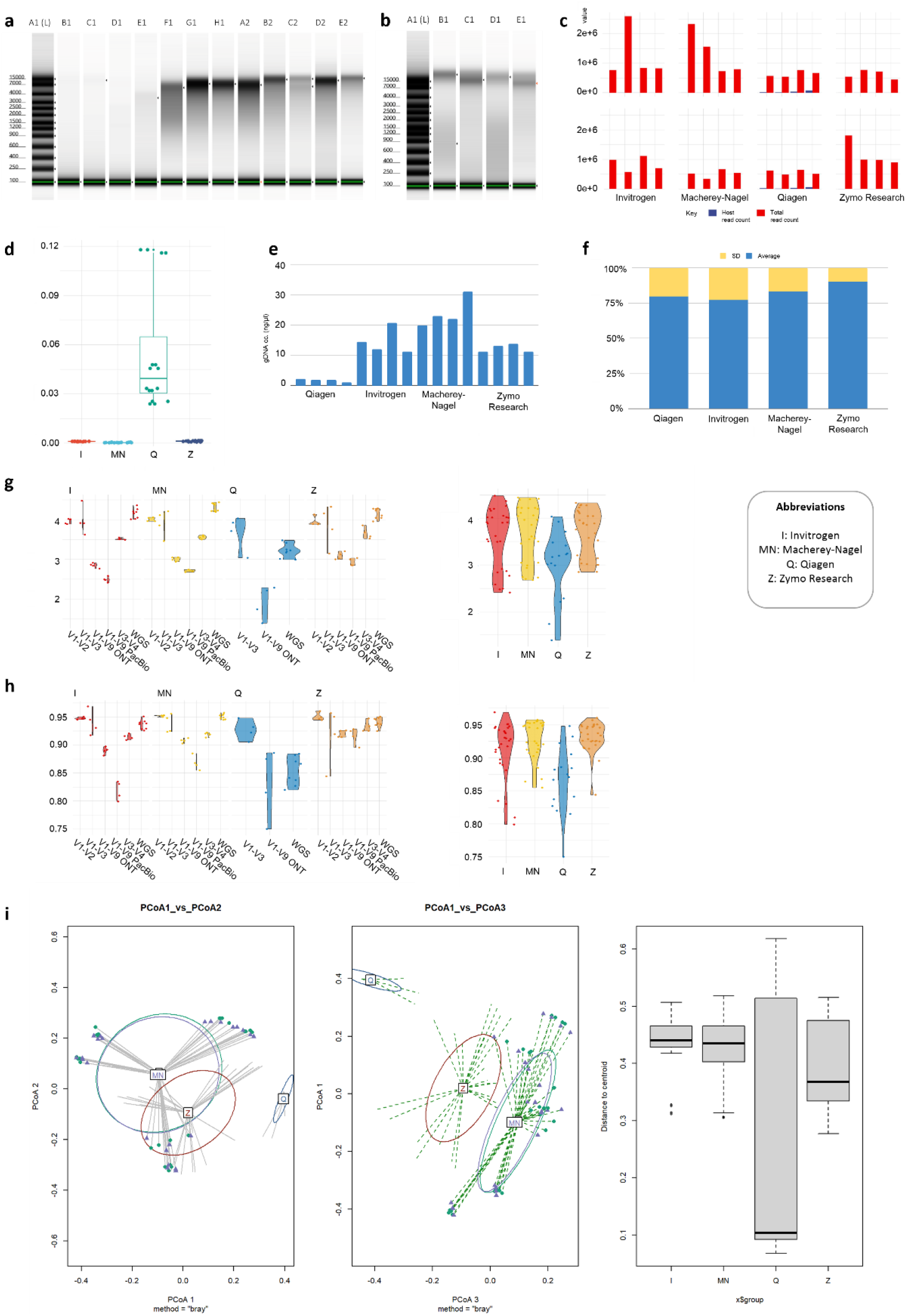
Impact of DNA isolation kit on yield, quality, and bacterial composition analysis. **a-d**, ***Quality of the Isolated DNA*** We observed significant differences in the molecular weight of DNA obtained using various kits. The Qiagen kit produced the most degraded DNA (**2a** – lanes B1-E1). We used the Quick-DNA HMW MagBead Kit from Zymo Research to extract High Molecular Weight (HMW) DNA, ensuring the quality necessary for sequencing the complete 16S rRNA gene with ONT sequencing. However, our findings suggest that HMW DNA is not essential for sequencing the entire V1-V9 region. The DNA quality and quantity obtained using the Macherey-Nagel (**2a** –lanes F1-A2) and Invitrogen **(2a** –lanes B2-E2) kits were suitable for long-read sequencing (LRS). The variation in average DNA fragment length produced by these three kits does not significantly affect V1-V9 sequencing (**2a** - lanes F1-E2 and **2b**). It’s important to note that for LRS-based whole-genome sequencing, maintaining HMW DNA integrity is essential. We found that the Zymo kit might be a better option for this purpose based on DNA length (**Supplementary Data 3**). Regarding DNA quality, we assessed fragment length and the kit’s efficiency in selectively extracting bacterial DNA. Even when using the “Isolation of DNA from Stool for Pathogen Detection” protocol with the Qiagen kit, the ratio of host DNA was significantly higher compared to the other three isolation kits (**2c-d**). **e-f**, ***Yield and Reproducibility of the Isolated DNA*** Among the four DNA extraction kits we tested on canine stool samples, all but the Qiagen kit produced DNA of adequate quantity and quality for sequencing (**2e-f**, **Supplementary Data 4a**). When we evaluated the Qiagen kit using stool samples from six different dogs, we consistently found below-average yields and quality (**Supplementary Data 4b, Supplementary Fig. 1**). Achieving consistent results from samples proved to be challenging when using the Qiagen kit, whereas the other three kits consistently delivered very similar quality for each replicate. Regarding concentrations, the Zymo kit showed the most consistent results, followed by Macherey-Nagel, with Qiagen exhibiting the most variation (**2e**-**f**). **g-i**, ***Microbial Composition, Diversity and Dispersion*** Alpha-diversity comparison between the different library preparation methods, according to each DNA isolation method: g, Shannon index; h, Simpson index (**g-h**). Beta-diversity analysis of each DNA isolation method was conducted. The PCoA plot, generated using the sample-wise Bray-Curtis distances, shows a large overlap between the Invitrogen and Macherey-Nagel methods, a partial overlap of Zymo, and Qiagen clustering farther from all other three methods. The associated boxplot displays the distances of each sample to its group centroid (**i**).

To test how significantly the DNA isolation kits influences the variation of the observed microbial composition, we used the same database and bioinformatic approach. Given that commonly used pipelines are applicable for particular sequencing methods, we used the minitax software, with a genome collection from NCBI as reference.

Previous studies demonstrated that α-diversity indices could successfully assess the efficiency of DNA extraction^17^. Here, we revealed a significant and consistent reduction in both richness (Shannon index) and evenness (Simpson index) using the Qiagen method compared three others (p-values < 0.000, for every comparison, using ANOVA) (**Figs. 2g-h**). Other pairwise richness comparisons were not significant. However, the Invitrogen kit showed lower evenness than the Zymo kit (**Supplementary Data 4**). We established a read number threshold for each sequencing approach where the α-diversity values remained stable and did not exhibit any significant reduction (**Supplementary Fig. 3**).

For a comprehensive evaluation of β-diversity, a sample-wise Bray-Curtis distance matrix was calculated. Subsequent analyses, including Non-Metric Multi-Dimensional Scaling (NMDS), Permutational Analysis of Dispersion (PERMDISP), and Permutational Analysis of Variance (PERMANOVA), were carried out using these distance values. This revealed distinct compositional differences attributed to the choice of DNA isolation method. The full-model PERMANOVA revealed that the DNA isolation had a substantial impact across various sequencing methods (WGS or 16S-Seq) and platforms. It accounted for roughly 28.2% of the overall variation observed in the microbial communities (p < 0.001).

An examination of sample dispersions for each DNA isolation method across all libraries, represented by the mean distance from the centroid in multivariate space (as estimated by PERMDISP2), revealed discernible variability. Such differences in dispersion can significantly influence the outcome of community composition analysis. These differences and similarities are graphically depicted in the Principal Coordinate Analyses (PCoA) for each isolation method (**Fig. 2i**).

Our dispersion analyses, carried out separately for each library, disclosed that Invitrogen consistently exhibits both the lowest average dispersions and the smallest standard deviations, with a mean distance to centroid of 0.0626 and a standard deviation of 0.0586 (**Supplementary Fig. 4a**). The Macherey-Nagel kit also yields low dispersions across almost every library, except the V1-V3. Compared to these two methods, the Zymo kit displayed results with increased dispersion in the LRS libraries and comparable dispersions in the SRS-based methods. The dispersion of the results obtained using Qiagen kit was comparable to other methods in the two libraries suitable for in-depth analysis (Illumina V1-V3 and WGS). Subsequent PERMANOVA evaluations for each library preparation method highlighted the influence of the DNA isolation techniques across different V-regions (and WGS) and systems. These results showed that the Macherey-Nagel and Invitrogen DNA isolation methods consistently produced microbial community compositions that were similar across the range of library preparation protocols. In contrast, the results of the Zymo and Qiagen methods showed community compositions that diverged significantly from those of Macherey-Nagel and Invitrogen, and often from each other as well. These are graphically depicted in the PCoA for each library calculated from the library-wise PERMIDISP results (**Supplementary Fig. 4b**). The pairwise comparisons further underscored these distinctions, with significant differences evident across the entire V-regions. Notably, the only exception to this trend was the Invitrogen/Macherey-Nagel pair.

### 3. Assessment of library construction and sequencing methods

In this part of the study, we aimed to elucidate the consistency or variability among the six library preparation methods. To achieve this, we compared the results of each DNA isolation technique for each sequencing library. Due to quality and quantity concerns with the Qiagen kit, only the WGS, V1-V3, and ONT V1-V9 libraries were prepared from these samples.

#### 3.1. Quality, Yield and Reproducibility

The ONT 16S library preparation demands the least experimental effort, whereas the Illumina WGS and PacBio 16S library preparation necessitate the most hands-on time (**Supplementary Fig. 5**). Apart from the PerkinElmer method, which often resulted in two nonspecific DNA fragments (400 and 800 bps), the other kits demonstrated excellent performance in terms of quality, yield, and consistency for dog samples (**Supplementary Fig. 6**). Yet, Nanopore technology falls somewhat behind in read quality (**Supplementary Data 5**).

#### 3.2. Microbial Composition, Diversity and Dispersion

The α-diversity analysis showed that in the SRS libraries, both richness and evenness measures were comparable among the three kits, with the exception of Qiagen. In the LRS libraries, Invitrogen showed somewhat lower diversity, Zymo the highest, while Macherey-Nagel also showed similarly high values.

We conducted β-diversity analyses using sample-wise Bray-Curtis distances. The NMDS plot depicted in **Fig. 3a** illustrates that the samples were more inclined to cluster according to their library preparation protocols than the DNA isolation methods employed. Indeed, in a full-model PERMANOVA, library selection was identified as the dominant factor in determining microbial community structures, accounting for 58.8% of the total observed variation. In addition to PERMANOVA, we also carried out PERMDISP analysis for each sample (**Fig. 3b**). The PCoA visualization indicated that no DNA isolation method produced the same microbial compositions across the various libraries. In fact, only the V1-V2 and V1-V3 libraries showed considerable overlaps in the case of Invitrogen and Macherey-Nagel (**Figs. 3b,c**). Additionally, the LRS and WGS libraries produced from Zymo DNA displayed results similar to those generated by Invitrogen and Macherey-Nagel. A heatmap from the β-diversity analysis depicted in **Fig. 3d** shows the pairwise comparisons for each combination of DNA isolation and library preparation method.

**Fig. 3:**
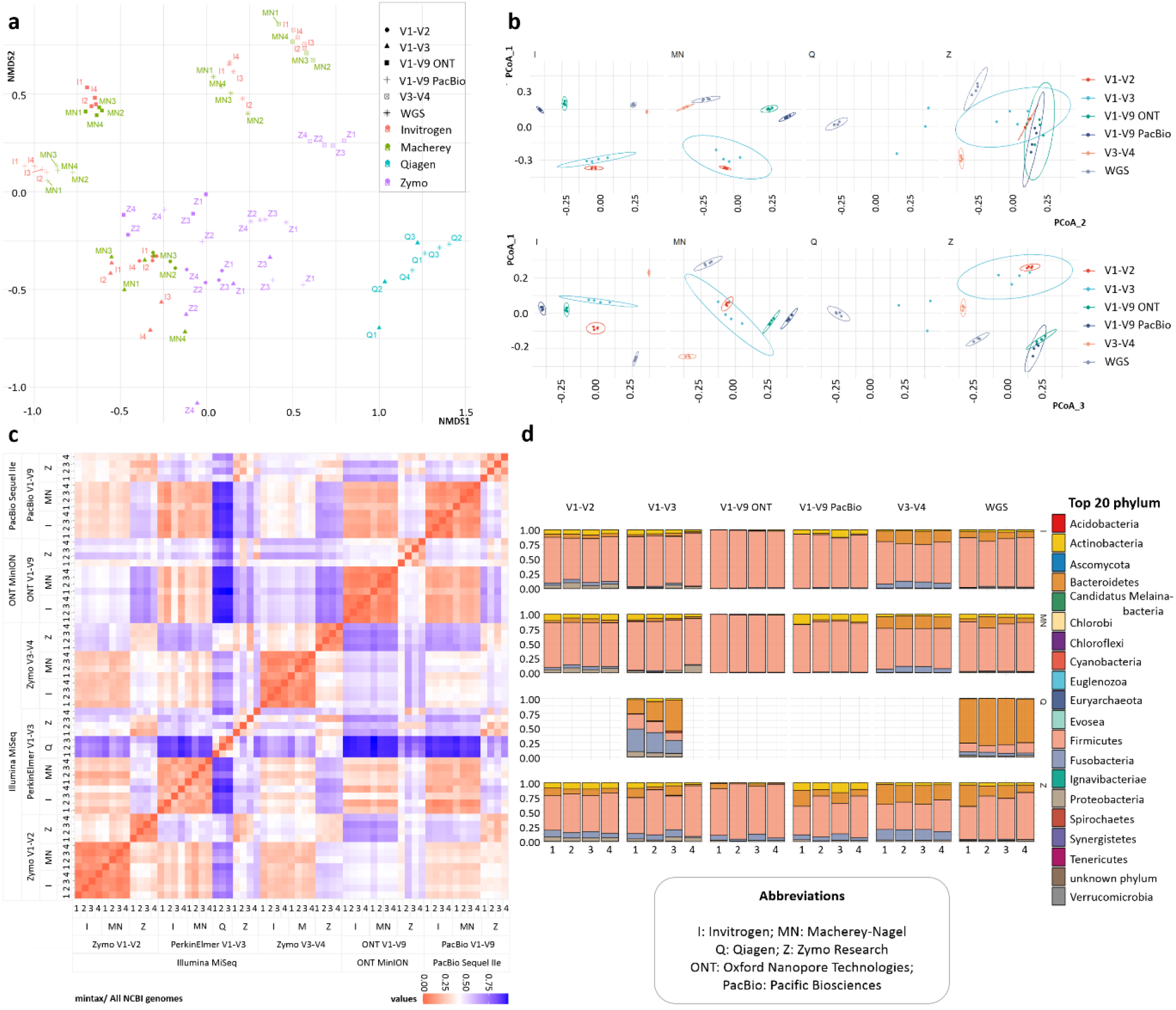
Comparison of library preparation protocols. **a,** In the NMDS plot, samples predominantly cluster based on their sequencing protocols, illustrating their dominant effect on microbial community compositions. There is noticeable spread in samples extracted using the Invitrogen and Macherey-Nagel DNA isolation methods, indicating variability in community compositions according to the library preparation protocol. Interestingly, the V3-V4 group clustered far from the V1-V3 group, which was evidently more similar to the V1-V2 group. The Zymo method showed somewhat closer grouping, while the WGS and V1-V3 samples from the Qiagen kit cluster together closely. Overall, the plot emphasizes the combined influence of both DNA isolation methods and sequencing protocols on community profiles, with sequencing protocols appearing to be the more influential factor. **b,** PCoA plots illustrating microbial composition variations across different DNA isolation methods and library preparations. The plots reveal that the Illumina V1-V2 and Illumina V1-V3 libraries have noticeable overlaps in the Invitrogen and Macherey-Nagel methods. The Zymo DNA extraction method displays significant overlaps between the ONT V1-V9, PacBio V1-V9, and Illumina WGS libraries. However, consistency within library preparation varies, as indicated by the spread of samples. For instance, the Illumina V3-V4 method in the Invitrogen quadrant shows tight clustering, suggesting more consistent results. **c,** The Bray-Curtis distance was calculated to compare the beta-diversity differences among sequencing data obtained from different sequencing technologies. Irrespective of the isolation kit used, the PacBio results are similar for both V1-V2 and V3-V4 regions (with the latter two being most alike). ONT and WGS show resemblance, especially ONT Zymo. Upon examining WGS, it is evident that Qiagen is distinct, only resembling itself, while the three other DNA isolators sho similarities with each other. For the V3-V4 and V1-V2 regions, as well as for PacBio, it is apparent that Invitrogen and Macherey-Nagel are more similar to each other than to Zymo. When comparing the libraries against themselves, it is evident that V1-V3 performs the least favorably. **d,** Our analysis showed that the usage of the V1-V3 library results in a higher proportion of the Firmicutes (86.43%), and in lower proportions of the phyla Bacteroidetes (2.69%) and Fusobacteria (2.28%), compared to the other two SRS-based 16S libraries. The V1-V2 and V3-V4 libraries showed a more balanced distribution, with a proportion of Bacteroidetes 10.86% and 12.37%, respectively and Fusobacteria 8.77% and 8.18%, respectively. In our V1-V9 analysis, we observed that the phylum Firmicutes dominates in both ONT and PacBio samples. However, its presence is even more pronounced in ONT samples, where it ranges from 96.24% to 99.19%, compared to PacBio samples, where it’s between 84.70% and 91.53%. Additionally, both methods barely detect Proteobacteria and greatly under-represent Fusobacteria (compared to the SRS methods) and Bacteroidetes among the top five phyla. The nanopore technique predominantly favored Firmicutes (89.28%), thereby constraining Actinobacteria and Bacteroidetes to low read counts as well. In contrast, PacBio showed a more balanced representation for these last three phyla (77.92% Firmicutes, 11.31% Actinobacteria, and 6.79% Bacteroidetes).

Since the overlaps in **Fig. 3b** were only partial and a noticeable difference in variability was observed across the different libraries, we assessed the distances of each sample from the centroid of its respective group (dispersion). We observed that the V1-V3 libraries displayed considerable variability in samples prepared using each DNA extraction method. With the exception of the V1-V3 and LRS libraries prepared with Zymo DNA, all other library preparation methods produced results exhibiting relatively low variability (**Supplementary Fig. 4c**).

Pairwise PERMANOVA, with library preparation technique as the sole variable, unambiguously demonstrated that all libraries held significant differences from each other, even after correcting the p-values for multiple testing. The overlaps observed in the PCoA plot (**Supplementary Fig. 4b**) were determined to be non-significant according to the PERMANOVA, which assesses differences between group centroids. In the case of the V1-V3 library, however, detected differences are not solely attributed to variations between centroids, as they are also influenced by differences in group dispersions.

We compared our results and data with those of other studies^43,50,51,52,53,54^ (**Supplementary Data 6**, **Supplementary Fig. 7**), all of which focus on a single aspect of the canine gut microbiome. A unanimous finding across all these publications, including our own dataset, suggests the predominant presence of five phyla. However, their abundance varies significantly, as shown in the **Supplementary Fig. 6**, underscoring the impact of the DNA isolation kit used on the observed microbial community composition.

#### 3.3 Difference in ratio of Gram(+) and Gram(-) bacteria

To test whether certain DNA extraction method favor Gram-negative bacteria over Gram-positives, we aggregated species abundances based on their cell-wall staining characteristics and conducted a similar analysis. We found that, indeed, the observed differences among DNA isolation methods could be significantly attributed to the varying resistance of cell walls to treatments. Specifically, only the Invitrogen and Macherey-Nagel methods produced microbial profiles with near-identical ratios of Gram-positive to Gram-negative bacteria. In stark contrast, all other pairwise comparisons highlighted pronounced disparities in these ratios. Concerning the differences among libraries, we observed significant disparities in multiple pairwise comparisons, with the V3-V4 region often being particularly notable (**Supplementary Fig. 8**).

### 4. Comparison of DNA Isolation Methods and Sequencing Libraries on a Synthetic MCS

To evaluate the accuracy of our DNA isolation methods and library preparation protocols, we employed a microbial community standard (MCS; Zymo D6300) of established composition as a reference standard (**Fig. 4b**, **Supplementary Fig. 9**, and **Supplementary Data 7**). Unlike the fecal DNA sample, where we evaluated the uniformity of results within each group of methods (extraction, library preparation and bioinformatics methods) and their similarity to one another, our primary goal in this context was to determine the ability of each group to accurately replicate the expected composition. Subsequently, we utilized the minitax software and the NCBI genome collection to compare the observed microbial compositions with the expected one.

**Fig. 4:**
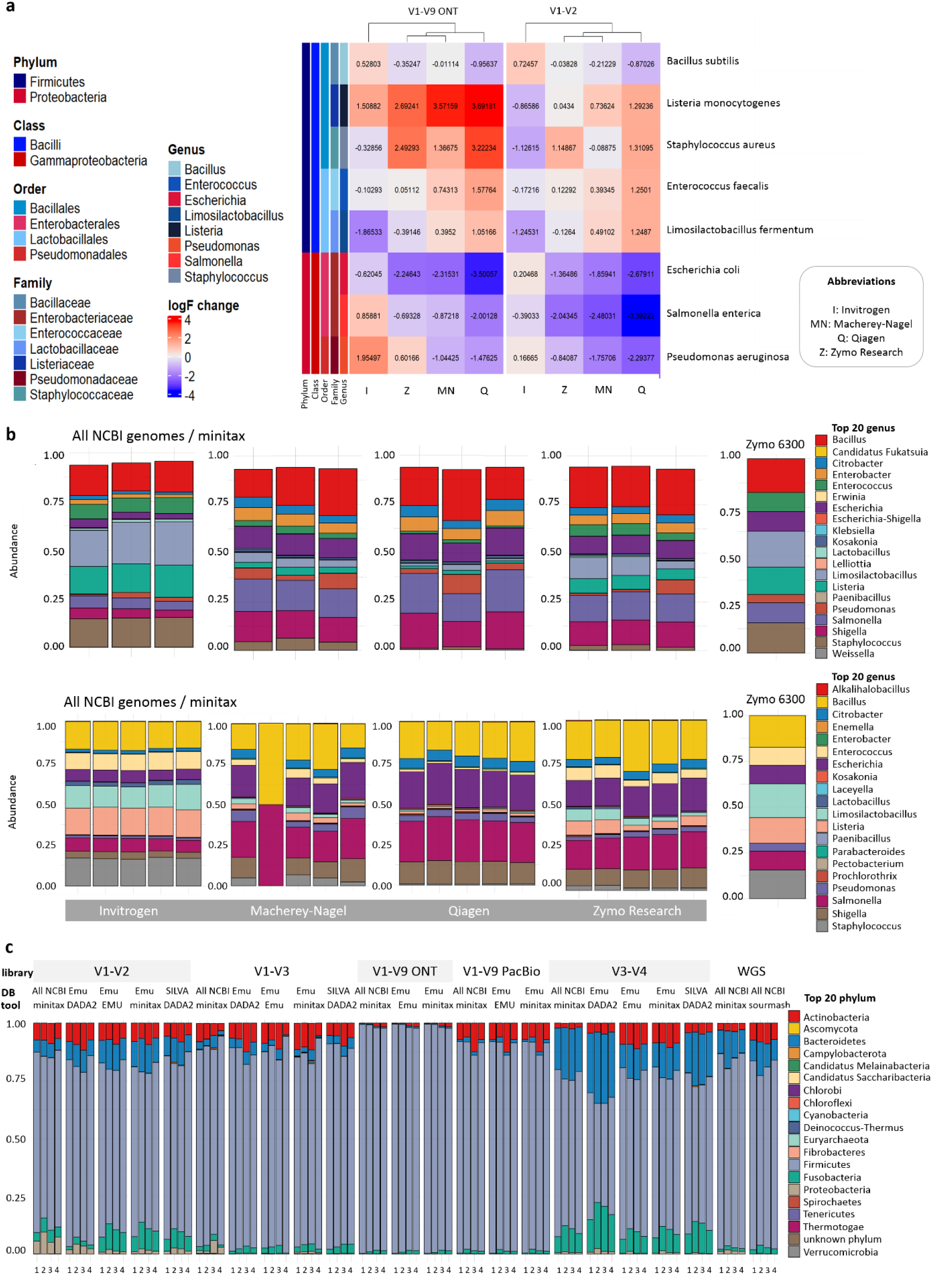
Comparative analysis of DNA isolation and library preparation protocols in MCS. **a,** We isolated DNA from the ZymoResearch MCS sample using the four isolation kits also employed for the dog samples. MCS is a mix of eight bacteria, of which five belong to the Firmicutes phylum and three are Proteobacteria. Subsequently, we generated Illumina V1-2 and ONT V1-9 libraries from the DNA samples. The abundance values identified by the Emu application were normalized to the theoretical values provided by Zymo and represented as log2 fold changes using the DESeq2 software. The data were visually represented in the heatmap. The study revealed that the data obtained using the Invitrogen kit showed the highest concordance with the theoretical values. **b,** Barplots showing the top 20 phyla in MCS samples using the Illumina V1-V2 and ONT V1-V9 methods, according to the DNA isolation methods, analyzed with minitax using the NCBI genome collection database. **c,** Barplot showing the top 20 phyla in the MCS samples, sequenced using Invitrogen DNA isolation kit, according to library preparation protocols, analyzed with minitax, DADA2 and Emu programs using several different databases.

#### 4.1 Microbial Composition, Diversity and Dispersion

The dispersion analysis showed that for the SRS library, in alignment with our observations from fecal DNA samples, the Invitrogen method exhibited the lowest dispersion. It is noteworthy that in the LRS library, the Qiagen method demonstrated similarly low variability. However, a contrasting scenario emerged upon examining the observed microbial compositions. Unlike the strong similarity observed *in vivo*, the Macherey-Nagel and Invitrogen methods showed significant divergence when evaluating the MCS. Although the Qiagen method remained the most distinct from Invitrogen, both Zymo and Macherey-Nagel exhibited a closer association with each other and a notable separation from Invitrogen. This trend was evident across both library types, underscoring the intrinsic differences between *in vivo* and *in vitro* sample evaluations (**Fig. 4c**).

#### 4.2. Difference in ratio of Gram(+) and Gram(-) bacteria

Qiagen tended to overrepresent Gram-negative bacteria in MCS samples and showed higher ratios for these bacteria in *in vivo* samples. Intriguingly, while the Macherey-Nagel method displayed a similar overrepresentation in fecal samples, it did not exhibit this trend in the MCS context. For both kits, the main underrepresented Gram-positive bacterial orders were Lactobacillus/Limosilactobacillus, whereas the majority of the overrepresented Gram-negative orders were Escherichia and Salmonella. We suspect that the Macherey-Nagel method may favor extracting DNA from Gram-negative bacteria, as evident in the controlled setting of MCS. This bias could be obscured in fecal samples due to their complexity, affecting the lysis of both Gram-positive and Gram-negative bacteria.

### 5. Comparison of bioinformatics techniques

To eliminate the variability in canine microbiome composition obtained by the different DNA extraction kits, we only used the Invitrogen DNA isolation method for comparison. We chose the NCBI genome collection as a reference to reduce disparities arising from database selection (**Fig. 3c**). Our analyses spanned the phylum, order, and genus taxonomic levels.

In this section of the study, we compared the effects of different databases (Emu, NCBI genomes and SILVA) and bioinformatic programs (Emu, minitax, DADA2 and sourmash) on the obtained canine microbiome compositions. In order to eliminate the variability in the results contributed to the DNA extraction method, we only used the Invitrogen DNA isolation kit in these comparisons. Using a full-model PERMANOVA we found that beyond the DNA isolation, library preparation and sequencing methods, the chosen bioinformatics approaches and databases notably influence the outcome. The last two factors accounted for 59.4% of the variation in microbial community composition, whereas the ’library’ factor accounted for only 20.1%, leaving 20.4% of the variation unexplained.Subsequent pairwise comparison showed that the differences among the software used with the Emu database were notably significant for DADA2 compared to the other two programs, while minitax closely resembled Emu. Using DADA2 with the SILVA database highlighted even starker differences. With minitax and the NCBI genome collection, differences from the Emu database results were significant only for the ONT V1-V9 library and from sourmash for WGS. However, consistent results were seen across all library types, regardless of the software/database pairing. Therefore, the minitax/NCBI genomes combination proved apt for evaluating DNA isolation kits and library types (**Fig. 4d**).

We also analyzed our MinION-sequenced samples using the ONT 16S Barcoding Kit alongside the 16S module of the EPI2ME Labs software, which performs a streamlined analysis process and quickly generates a Sankey tree diagram on its online platform. We found that EPI2ME produces discrepancies at the genus level (**Fig. 5**). To verify this finding, we extended our study beyond the original canine stool sample (**Fig 5.a,b**), incorporating fecal samples from six additional dogs (**Fig. 5c**), along with both neonatal^55^ (**Fig. 5d**) and adult^56^ (**Fig. 5e**) human stool specimens, adult human skin^57^ (**Fig. 5f**), and MCS (**Fig. 5g**).

**Fig. 5:**
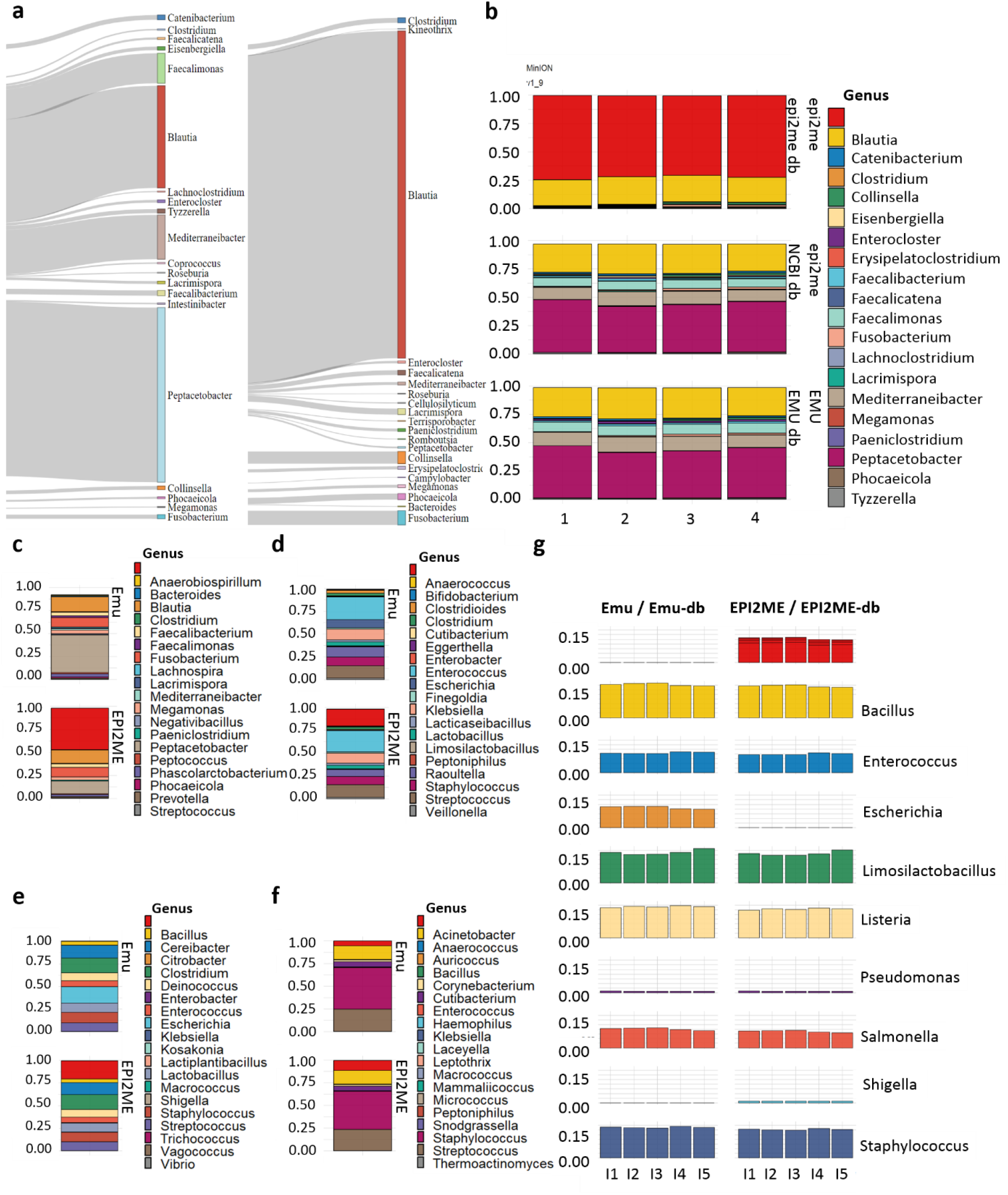
The EPI2ME analysis platform yields inaccurate genus-level data. **a-b,** The 16S module of the EPI2ME Labs program by ONT offers an easy-to-use analysis and quickly creates intuitive Sankey tree diagrams on its online platform. We ran our samples, sequenced on the MinION platform using the ONT 16S kit, and initially observed the Blautia genus to be the predominant component, with an average occurrence of about 80%. This result was consistent across all DNA isolation kits but contradicts both literature data and sequencing results from other methods (Illumina WGS, V1-V2, V3-V4, V1–V3, PacBio V1-V9). We also analyzed these samples using other programs (Emu and minitax). Our findings indicate that the EPI2ME program significantly enriches Blautia. However, when the same sequencing results are assessed with other software, the Blautia ratio aligns with the ∼15% reported in the literature^50^. Contrary to our initial assumptions, EPI2ME does not exaggerate the presence of Blautia but either excludes genera or underrepresents them. We analyzed the entirety of the ONT MinION V1-V9 datasets available in the databases using EPI2ME and compared the outcomes with those derived from Emu. In our canine samples, the software failed to detect any reads corresponding to the Peptacetobacter, Faecalimonas, and Mediterraneibacter genera, discarding 75% of the reads and skewing the actual compositional ratios. **a-b ,** The 16S module of the EPI2ME Labs program by ONT offers an easy-to-use analysis and quickly creates intuitive Sankey tree diagrams on its online platform. We ran our samples on the MinION platform using the ONT 16S kit and initially observed that the Blautia genus was the predominant component, with an average occurrence of about 80%. This result was consistent across all DNA isolation kits but contradicts both literature data and sequencing results from other methods (Illumina WGS, V1-V2, V3-V4, V1–V3, PacBio V1-V9). We also analyzed these samples using other programs (Emu and minitax). When the same sequencing results are assessed with other software, the Blautia ratio aligns with the ∼15% reported in the literature^50^. Contrary to our initial assumptions, EPI2ME does not exaggerate the presence of Blautia but either fails to identify some genera or underrepresents them. This was actually caused by the filtering out of other taxa based on the LCA tag. We analyzed the entirety of the ONT MinION V1-V9 datasets available in the databases using EPI2ME and compared the outcomes with those derived from Emu. In our canine samples, the software failed to detect any reads corresponding to the Peptacetobacter, Faecalimonas, and Mediterraneibacter genera, discarding 75% of the reads and skewing the actual compositional ratios. We reprocessed the data, substituting EPI2ME annotations with annotations based on NCBI taxon IDs. The outcomes, closely resembling those achieved with Emu, indicating that the LCA tag filtering in EPI2ME filtering may not be necessary. **c,** We sequenced fecal samples from six additional dogs to determine if this distortion at the genus level is conclusively due to bioinformatics. We performed sequencing of ONT V1-V9 libraries and analyzed the data using both EPI2ME and Emu. On average, EPI2ME discards 50% of the reads during the process from the six samples. The genus Peptacetobacter was largely undetected, and the genera Faecalimonas and Mediterraneibacter were underrepresented, similar to our reference sample. **d-f,** We downloaded the available independent ONT V1-V9 data from databases to compare EPI2ME and Emu. Accessible datasets included those from a neonate (**d**) and an adult human fecal (**e**) sample set, as well as a dataset from human skin (**f**) samples. For the fecal samples, EPI2ME does not analyze 15-25% of the data, and the genus Escherichia is entirely excluded. For skin samples, there is minimal variation between the analyses of the two programs, with approximately 5% of the reads not analyzed by EPI2ME, mostly belonging to the genera Cutibacterium and Staphylococcus. **g,** In the analysis of the microbial mix sourced from Zymo, we observed that EPI2ME did not include around 15% of the reads, and it failed to identify the genus Escherichia. Conversely, EPI2ME identified a genus (Shigella) that is not present in the mix.

We ascertained that the EPI2ME software excluded a substantial fraction of data during its processing due to the application of Lowest Common Ancestor (LCA) tags. Specifically, within the BLAST analysis module, the LCA tag was designed to filter out reads when the top three predicted genera do not match the tag. We reprocessed the EPI2ME data and, using the updated information, we assigned the reads to the NCBI database according to their taxon ID. This revealed a similarity in the results produced by EPI2ME and Emu. Therefore, employing the LCA tag to filter out data in EPI2ME is unnecessary and, in fact, diminishes the accuracy of the program (**Fig. 5b**).

### 6. Evaluation of minitax: Benchmarking Across Various Sequencing Method

For consistent comparisons across varied sequencing data, we developed minitax, a tool for metagenome sequencing taxonomy. We rigorously tested minitax across platforms and diverse datasets. We compared it against several bioinformatic programs mainly in terms of correlation between the observed and theoretical composition (as explained with r^2^ values), that is their ability to correctly reconstruct the reference microbial composition.

#### 6.1. Comparing minitax with Emu using ONT V1-V9 Sequencing of MCS

We employed the ONT platform to sequence the V1-V9 regions of the 16S rRNA gene from the MCS (Zymo D6300). We contrasted the performance of minitax with Emu, which, like minitax, utilizes minimap2 for read alignment. However, the expectation-maximization approach of Emu is not suited for WGS datasets. We found that the Invitrogen DNA isolation method consistently outperforms other methods (**Fig. 6a** and **Supplementary Fig. 10a**). When comparing the workflows, minitax and Emu delivered similar results when utilizing the Emu database, indicating the robustness of minitax. Notably, even though complete genome databases are not typically used for 16S-Seq, minitax efficiently utilizes them, providing solid reconstructions up to the genus level. This adaptability is beneficial for those wanting a consistent database strategy for both WGS and 16S sequencing. However, species-level precision using the NCBI database shows a substantial decline, underscoring the importance of choosing a database that aligns with the desired level of resolution.

**Fig. 6:**
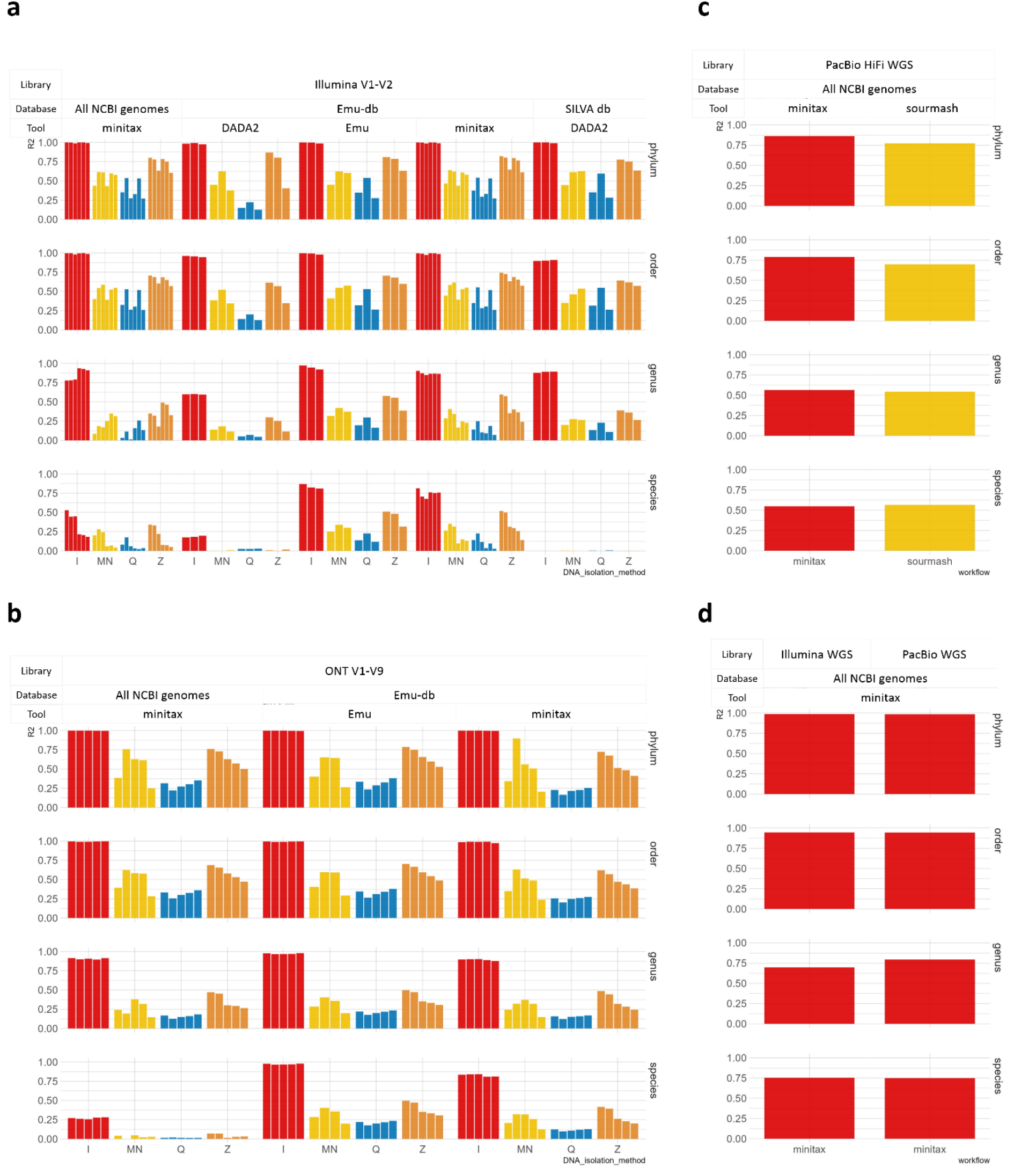
Comparative analysis of minitax performance across different sequencing platforms and datasets. **a, *ONT V1-V9 Sequencing: performance comparison of minitax against Emu for the Zymo D6300 microbial community reference*** We compared minitax to both Emu, using Emu-database, and DADA2, a popular tool designed for Illumina 16S amplicon sequencing, using both Emu-database and its standard SILVA database. The results highlight the exceptional performance of the Invitrogen method among various DNA isolation techniques. Notably, minitax remains impressively competitive against both Emu and DADA2, demonstrating its flexibility across 16S and WGS datasets. **b, *Illumina V1-V2 sequencing: minitax benchmarking against Emu and DADA2 for Zymo D6300 MCS*** Benchmarking of minitax against Emu and DADA2 for the Zymo D6300 MCS. We compared minitax to both Emu, using Emu-database, and DADA2, a popular tool designed for Illumina 16S amplicon sequencing using Emu-database and its standard SILVA database. The results underscore the outstanding performance of the Invitrogen method among various DNA isolation techniques. Notably, minitax remains impressively competitive against both Emu and DADA2, demonstrating its flexibility across 16S and WGS datasets. *c, PacBio HiFi WGS: minitax vs. sourmash performance comparison on Zymo D6331 MCS dataset* This figure contrasts the performance minitax with that of sourmash using a Zymo D6331 MCS dataset sequenced on a PacBio HiFi platform. We compared their performance with and without the unclassified reads, which contribute to the total community composition, thereby skewing the ratios for the identified taxa. When including unclassified reads, minitax shows significantly higher r2 values compared to sourmash on all taxonomic levels (0.430 and 0.555 for sourmash and minitax at the species level, respectively). However, when we exclude these reads, these differences decrease, and sourmash provides slightly higher values at the species level (0.548 vs 0.566). **d, *CAMISIM mouse gut dataset: comparative analysis of minitax performance on Illumina and PacBio platforms*** Up to the family level, minitax demonstrated very strong, near-identical correlations between the theoretical and observed taxonomic compositions: r2 of 0.9547 and 0.9515 for Illumina and PacBio, respectively. However, the genus-level classifications revealed a noticeable disparity between the two platforms. While Illumina reached an r2 of 0.4417, PacBio demonstrated a markedly higher r2 of 0.5892. At the species-level identification, the r2 values were 0.2128 for Illumina and 0.3983 for PacBio, with PacBio showing superior performance compared to Illumina.

#### 6.2. Comparing minitax with Emu and DADA2 using Illumina V1-V2 Sequencing of MCS

In this work, we utilized an Illumina-sequenced V1-V2 dataset derived from our MCS (Zymo D6300) DNA sample. The results consistently showed that the Invitrogen method outperforms other methods (**Fig. 6b** and **Supplementary Fig. 10b**). In terms of software performance, both Emu and minitax workflows markedly outperform DADA2, even when using the standard SILVA database, which is commonly utilized by DADA2. It is particularly noteworthy that while Emu and DADA2 are optimized for amplicon sequencing, minitax remains competitive, especially given its versatility in managing both 16S and WGS datasets.

#### 6.3. Comparing minitax with sourmash using MCS data of PacBio HiFi WGS

We compared minitax to sourmash, the latter being acknowledged as one of the leading software choices for both LRS and SRS mWGS data^10^, using the PacBio HiFi dataset (NCBI accession: SRX9569057) from MCS (Zymo D6331). We carried out the comparison of r^2^ values with and without excluding unclassified reads. Minitax outperformed sourmash when excluding unclassified reads, yet fell marginally short at the species level when these reads were included (**Fig. 6c** and **Supplementary Fig. 10c**), as this distorts the relative abundances of the identified taxa. These results underscore the effectiveness of minitax in handling such datasets.

#### 6.4. CAMISIM: Simulated Mouse Gut Datasets

Expanding our validation range, we examined simulated datasets from the CAMISIM mouse gut project, consisting of 10 samples each from PacBio and Illumina. For both data types minitax exhibited reliable performance, reaching an r^2^=0.96 at the phylum level, which decreased to r^2^=0.59 at the species level (**Fig. 6d**).

### DISCUSSION

In this comprehensive study, we compared metagenomic methods using canine fecal samples and a synthetic microbial community standard. The primary objective was to interrogate the fidelity of various methodological approaches including DNA extraction, library preparation, and bioinformatics analysis. To facilitate this, we developed ’minitax’, a versatile bioinformatics tool designed for use with diverse metagenomic laboratory protocols.

Choosing the right DNA isolation method is vital for maximizing yield and reducing fragmentation, thus preserving the integrity of further analysis^17^. The best extraction kit should deliver high-quality, high-yield DNA to reduce the risk of false negatives^58^; remove PCR inhibitors found in fecal samples^59,60,61,62,63^, provide consistent results, and effectively break down Gram-positive bacterial cell walls for a true reflection of community composition^9,24,25,26,27^.

In our work, we assessed the effectiveness of multiple DNA extraction techniques in terms of the quality and quantity of extracted DNA, and determined whether they produce accurate representations of the microbial composition. The Invitrogen and Macherey-Nagel methods emerged as particularly robust, yielding consistent microbial profiles across various library preparations. In stark contrast, methods from Zymo and Qiagen introduced marked biases, demonstrating the profound impact of DNA extraction can have on downstream analyses.

Recent studies^12,20^, including our own, indicates that WGS offers greater taxonomic diversity than 16S-Seq, however, unlike SRS platforms targeting 16S V-regions, full 16S gene sequencing significantly improves bacterial community identification. We extended our analysis to include both SRS and LRS platforms, as well as 16S rRNA gene amplicon sequencing and mWGS strategies. We observed that the V1-V3 libraries showed significant variability across all DNA extraction techniques, especially with the Zymo kit, and to a lesser degree with the Invitrogen and Macherey-Nagel methods. However, when excluding the V1-V3 libraries and the LRS libraries created using the Zymo DNA, the rest of the library preparation techniques exhibited minimal variation. Notably, this variability was observed irrespective of the DNA extraction kit employed, reaffirming the necessity for careful selection of library preparation methods to ensure accurate microbial community profiling. Next, we carried out an in-depth assessment of 16S V1-V9 sequencing using both ONT and PacBio platforms. Notably, only the Zymo method produced consistent results across both libraries. Additionally, we assessed both a V1-V2 and a V1-V9 library on MCS. We obtained that the Invitrogen method yielded the closest match between the actual and ascertained compositions in both libraries. The Macherey-Nagel method was more similar to the approach of Zymo and significantly differed from Invitrogen in both SRS and LRS libraries. These findings underscore the varying performances between *in vitro* and *in vivo* sample analyses, thereby highlighting the inapplicability of the MCS for validation purposes in other systems.

The significance of the applied bioinformatic tool in metagenomic analysis has been discussed by others^33^. We also addressed this issue and found that approximately 60% of the variation in microbial community profiles is attributed to computational factors. Both Emu and minitax consistently yielded results that were closely aligned, and for MCS samples, they correlated highly with the true compositions of each amplicon library. A similar pattern was evident between sourmash and minitax for the mWGS library, underlining the versatility of the minitax tool. In contrast, the results obtained by DADA2 pipeline varied considerably from those of minitax and Emu in both *in vivo* and *in vitro* samples.

Finally, we tested our bioinformatic tool across a variety of sequencing platforms and libraries, contrasting it with well-regarded and efficient tools like Emu and sourmash. Although minitax did not always yield the highest correlation values in every instance, it frequently outperformed these methods, delivering consistent and reliable results. Due to its comprehensive database, the results from minitax can be compared between mWGS and amplicon-based libraries.

In summary, our research offers an in-depth assessment of the gut microbiome, underscoring that methodological optimization and uniformity are important for ensuring accurate representation and reproducibility. However, it is important to note that expanding the range of methodologies can also be advantageous as it helps to surmount the intrinsic limitations of a given approach, thereby enabling a more thorough understanding of the subject under investigation^2^.

### ONLINE METHODS

#### SAMPLE COLLECTION

A fecal sample from a 13.5-year-old healthy male Pumi (a Hungarian purebred dog) was collected within 1 minute of defecation and was immediately frozen and stored at −80°C. Stool samples from six other healthy Pumi dogs (four puppies, a 7-year-old female, and a 6.5-year-old sterilized male) were used as controls (**Table 1**). Furthermore, the ZymoBIOMICS Microbial Community Standard (MCS; Zymo Research, D6300) was used to validate the obtained data.

**Table 1.** Summary table of basic characteristics of the dogs enrolled in this study. * sterilized; underlined names are used in data identifiers in ENA.

| Sample ID in figures |  | Age | Registered name 'call name' | Sampling time | Living place at sampling time |
| --- | --- | --- | --- | --- | --- |
| Number | Gender |  |  |  |  |
| - (main subject of the study) | Male | 13,5 years | Harsányi-Tekergő Bátor 'Toto' (<u>Toti</u>) | Nov 20, 2021 | Szeged, Hungary |
| 1 | Female | 7 years | Galla-Hegyi Suba 'Sugó' | May 08, 2021 | Baja, Hungary |
| 1 | Male * | 6,5 years | Vöröskői Kondacsipkedő Lidérc 'Loki' | May 13, 2021 | Baja, Hungary |
| 2 | Female | 6 months | Duna-menti Dumás <u>Boszi</u> | May 9, 2021 | Baja, Hungary |
| 3 | Female | 6 months | Duna-menti Dumás <u>Borcsa</u> | May 4, 2021 | Budapest, Hungary |
| 2 | Male | 6 months | Duna-menti Dumás <u>Blöff</u> | May 5, 2021 | Budakalász, Hungary |
| 3 | Male | 6 months | Duna-menti Dumás <u>Briós</u> | May 4, 2021 | La Tour-de-Peilz, Switzerland |

#### DNA PURIFICATION

In our work, the following commercially available DNA isolation kits were tested using four technical replicates for each: 1. QIAGEN QIAamp Fast DNA Stool Mini Kit; 2. Invitrogen PureLink™ Microbiome DNA Purification Kit; 3. Macherey-Nagel NucleoSpin DNA Stool Mini kit; and 4. Zymo Research Quick-DNA™ HMW MagBead Kit. DNA extraction was conducted based on the guidelines supplied with the kit. A brief overview of their procedures is provided below.

#### QIAGEN QIAamp Fast DNA Stool Mini Kit

##### Canine stool

A 200 mg stool sample was placed in a 2 ml microcentrifuge tube and kept on ice. InhibitEX Buffer was added to each sample, and they were mixed using a vortex until completely homogenized. The large stool pieces were removed by centrifugation. Six hundred μl of the supernatant was combined with 25 μl of proteinase K and 600 μl of Buffer AL. The mixture was thoroughly vortexed, then heated to 95°C for 5 min (this is a 95°C lysis incubation for 5 minutes, diverging from the 70°C recommended by the QIAamp kit). Six hundred μl of 100% ethanol was added to the lysate and then mixed. Six hundred μl of the lysate was loaded onto a QIAamp spin column and centrifuged at 20,000 x g for 1 min. The QIAamp spin column was placed into a new 2 ml tube. The remainder of the lysate was then loaded onto the column. After centrifugation, 500 μl of Buffer AW1 was added to the column. This was followed by a 20,000 x g centrifugation for 1 min, then we discarded the collection tube. Next, 500 μl of Buffer AW2 was added to the column, which was then placed into a new collection tube. A full-speed centrifugation (20,000 x g) was performed for 3 min. To avoid any Buffer AW2 carryover, the spin column was set in a fresh 2 ml collection tube and the samples were spun down at full speed for 3 min. The spin columns were then transferred to new Eppendorf tubes, and 100 μl of Buffer ATE was directly loaded onto the QIAamp membrane. After letting it incubate at room temperature for 1 min, a centrifugation (20,000 x g) step was carried out for 1 min to elute the DNA in 50 μl in elution buffer, followed by storing the DNA solution at -20°C.

##### MCS sample

75μl from the mix was used for DNA purification, following the same protocol as for the dog sample, with the DNA being eluted in a final volume of 50 μl

##### Invitrogen PureLink™ Microbiome DNA Purification Kit

###### Canine stool

A 200 mg sample was combined with 600 μl of S1 Lysis Buffer in the Bead Tube (provided in the kit) and the mixture was homogenized by vortexing. Subsequently the 100 µL of S2 Lysis Enhancer (from the kit) was added, and the samples were vortexed again. The mixtures were then incubated at 65°C for 10 minutes. For homogenization, the samples were subjected to bead beating using a vortex mixer with horizontal agitation at maximum speed for 10 min. The samples were then centrifuged at 14,000 × g for 5 min. Afterwards, 400 µL of the supernatant was transferred to a new Eppendorf tube and mixed with 250 µL of S3 Cleanup Buffer. The samples were centrifuged again at 14,000 × g for 2 min, and 500 µL of the resulting supernatant was transferred to a clean tube and mixed with 900 µL of S4 Binding Buffer. After brief vortexing, 700 µL of the sample mixture was loaded onto a spin column and centrifuged at 14,000 × g for 1 min. The spin column was then placed in a new tube, and the remaining sample mixture was loaded onto it for an additional 1 min centrifugation. The spin column was subsequently placed in a clean collection tube, and 500 µL of S5 Wash Buffer was added, followed by centrifugation at 14,000 × g for 1 min. To remove any residual S5 Wash Buffer, a second centrifugation was carried out at 14,000 × g for 30 sec. Finally, DNA was eluted from the spin columns using 100 µL of S6 Elution Buffer and stored at -20°C.

###### MCS sample

75 μl of the microbial mix was utilized for DNA extraction, following the same steps as described above. The DNA was eluted in 50 μl S6 Elution Buffer.

##### Macherey-Nagel NucleoSpin DNA Stool Mini kit

**Canine stool**: DNA isolation was performed using 200 mg of fecal sample, which was transferred to a Macherey-Nagel Bead Tube Type A, and then 850 μL Buffer ST1 was added. The mixtures were shaken horizontally for 3 seconds before being placed in a heat incubator. Subsequently, the samples were incubated for 5 min at 70 °C, agitated on a Vortex-Genie® 2 at full speed and room temperature for 10 min, and then centrifuged for 3 min at 13,000 x g. Six hundred μl of the supernatant was transferred to a new 2 ml tube, and 100 μl of Buffer ST2 was added and briefly vortexed. The mixtures were incubated for 5 min at 4 °C and then centrifuged for 3 min at 13,000 x g. Five hundred fifty μl of the lysate was loaded onto a NucleoSpin® Inhibitor Removal Column and centrifuged for 1 min at 13,000 x g. The Inhibitor Removal Column was discarded. Two hundred μl of Buffer ST3 was added to the samples, which were then mixed. Seven hundred μl of the sample mixture was loaded onto a NucleoSpin® DNA Stool Column and centrifuged for 1 min at 13,000 x g. The column was placed in a new tube. The sample was washed four times: first, 600 μl of Buffer ST3 was added to the NucleoSpin® DNA Stool Column and centrifuged for 1 min at 13,000 x g. The column was then placed in a new tube, and 550 μl of Buffer ST4 was added. After a 1 min centrifugation at 13,000 x g, the column was placed into a new tube, and 700 μl of Buffer ST5 was added. Following a brief vortexing, the samples were centrifuged for 1 min at 13,000 x g. The column was placed in a new tube, and 700 μl of Buffer ST5 was added, followed by a 1 min centrifugation at 13,000 x g. The flow-through was discarded, and the column was placed back onto the tube. The silica membrane of the column was dried by a 2 min centrifugation at 13,000 x g. One hundred μl of Buffer SE was loaded onto the center of the column, and the DNA was eluted by centrifugation for 1 min at 13,000 x g. DNA samples were stored at -20°C.

###### MCS sample

75 μl of the MCS mixture was utilized for DNA isolation, with the sample being eluted in a final volume of 50 μl.

##### Zymo Research Quick-DNA™ HMW MagBead Kit

**Canine stool**: One hundred mg of the fecal sample was used as initial weight and resuspended in 200 µl of DNA/RNA Shield™, followed by incubation at room temperature (20-30°C) on a tube rotator for 5 minutes. Subsequently, 33 µl of MagBinding Beads were added to each sample, mixed and placed on a shaker for a 10 min. The sample was then placed on a magnetic stand until a clear separation of the beads and the solution was observed, after which the supernatant was removed. For the washing step, 500 µl of Quick-DNA™ MagBinding Buffer was added and the beads were resuspended and shaken for 5 min. The sample was returned to the magnetic stand, and the supernatant was discarded. Next, 500 µl of DNA Pre-Wash Buffer was added, and the beads were resuspended. The sample was placed on the magnetic stand again, and the supernatant was discarded. In the subsequent step, 900 µl of g-DNA Wash Buffer was added and mixed, and the entire liquid was transferred to a new tube. The magnetic stand was used to separate the beads from the solution, and the supernatant was discarded. These washing steps were repeated once more. The sample were left to air dry for 20 min. For the elution step, 50 µl of DNA Elution Buffer was added to each sample. After mixing, the solution was incubated at room temperature for 5 min. Finally, the sample was placed back on the magnetic stand until the beads separated from the solution. The eluted DNA was carefully transferred to a new tube and stored at -20°C for future use.

**MCS sample**: 75 μl of the mix was used for DNA purification, following the same protocol as described for the dog sample. The DNA was eluted in a final volume of 50 μl.

##### LIBRARY PREPARATION

###### Zymo Research V1-V2

**DNA from canine stool**: Ten µl of Quick-16S™ qPCR Premix were mixed with 4 µl of Quick-16S™ Primer Set V1-V2 and 4 µl of ZymoBIOMICS® DNase/RNase Free Water. Additionally, 2 µl of DNA samples (2.5 ng/µl) were added. PCR was conducted in a Verity Thermal Cycler (Applied Biosystems) as per the Zymo Research Manual (**Table 2**). After amplification, 1 µl of Reaction Clean-up Solution was added to the samples, which were then incubated at 37°C for 15 min. The reactions were terminated by heating to 95°C for 10 min, and the samples were subsequently cooled to 4°C. Next, 10 µl of Quick-16S™ qPCR Premix and 4 µl of ZymoBIOMICS® DNase/RNase Free Water were combined. Index primers (2 µl each from ZA5 and ZA7, see **Table 3** for detailed pairs and sequences) and 2 µl of the amplified DNA were also measured into the mixture. Barcoded PCR reactions were performed as recommended by the manual (**Table 4**). For purification of the PCR products, Select-a-Size MagBeads were used. First, the MagBeads were resuspended by shaking, and then 16 µl of the Select-a-Size MagBeads were mixed with each sample. The mixture was incubated at room temperature for 5 min and then placed on a magnetic rack for 3 to10 min. The supernatant was discarded and the beads were washed twice with 200 μl of DNA Wash Buffer. The samples were removed from the magnet and were incubated for 3 min at room temperature to eliminate all traces of buffer. Libraries were eluted in 25 µl of DNA Elution Buffer and stored at -20°C until further use.

**Table 2.**
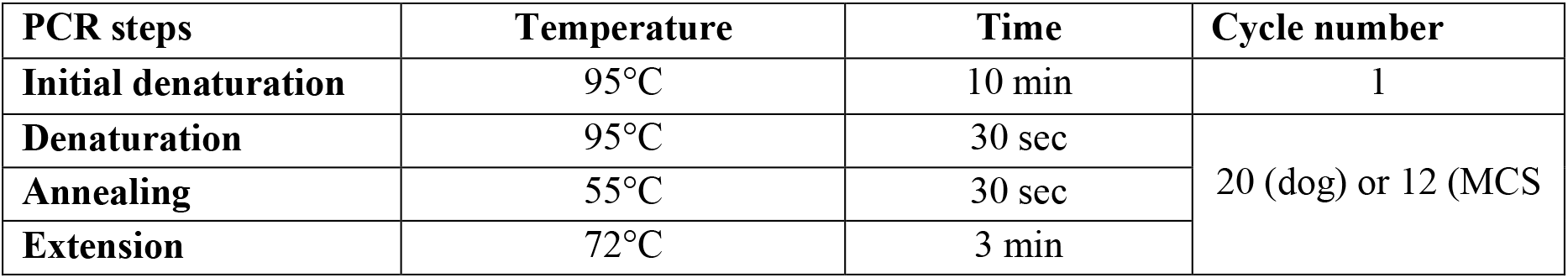
PCR amplification conditions for Zymo Research Quick-16S library preparation (V1-V2 and V3-V4 amplicon sequencing) – PCR I.

**Table 3.** Index barcode (ZA7-ZA5) combinations used for library preparation using the Zymo Research Quick-16S NGS Library Prep Kit. a, Barcodes used for canine stool samples; **b,** Barcode sequences applied for MCS samples Abbreviations: I: Invitrogen; MN: Macherey-Nagel; Z: Zymo Research.

| **a,** |  |  |  |  |
| --- | --- | --- | --- | --- |
| Sample | Index Barcode ZA7 | Sequence ZA7 | Index Barcode ZA5 | Sequence ZA5 |
| **I1** | ZA701 | ACCTGGAT | ZA501 | TTCTAGAC |
| **I2** | ZA702 | GTGCCATA | ZA502 | CCGATCTT |
| **I3** | ZA703 | TGAATCCG | ZA503 | TAAGATCC |
| **I4** | ZA704 | CATGATGC | ZA504 | AGGTCATT |
| **MN1** | ZA705 | AATGTCCT | ZA501 | TTCTAGAC |
| **MN2** | ZA706 | ATAGGCTC | ZA502 | CCGATCTT |
| **MN3** | ZA701 | ACCTGGAT | ZA503 | TAAGATCC |
| **MN4** | ZA702 | GTGCCATA | ZA504 | AGGTCATT |
| **Z1** | ZA703 | TGAATCCG | ZA501 | TTCTAGAC |
| **Z2** | ZA704 | CATGATGC | ZA502 | CCGATCTT |
| **Z3** | ZA705 | AATGTCCT | ZA503 | TAAGATCC |
| **Z4** | ZA706 | ATAGGCTC | ZA504 | AGGTCATT |
| **b,** |  |  |  |  |
| Sample | Index Barcode ZA7 | Sequence ZA7 | Index Barcode ZA5 | Sequence ZA5 |
| **I1** | ZA707 | TTGCGGAG | ZA505 | GTGTGTCA |
| **I2** | ZA705 | AATGTCCT | ZA504 | AGGTCATT |
| **I3** | ZA701 | ACCTGGAT | ZA501 | TTCTAGAC |
| **Q1** | ZA705 | AATGTCCT | ZA503 | TAAGATCC |
| **Q2** | ZA702 | GTGCCATA | ZA502 | CCGATCTT |
| **Q3** | ZA706 | ATAGGCTC | ZA504 | AGGTCATT |
| **MN1** | ZA703 | TGAATCCG | ZA501 | TTCTAGAC |
| **MN3** | ZA708 | GCCTTCCA | ZA508 | CCACAGGT |
| **MN4** | ZA704 | CATGATGC | ZA502 | CCGATCTT |
| **Z1** | ZA706 | ATAGGCTC | ZA505 | GTGTGTCA |
| **Z2** | ZA709 | GTCAGTCT | ZA508 | CCACAGGT |
| **Z3** | ZA705 | AATGTCCT | ZA505 | GTGTGTCA |

**Table 4.**
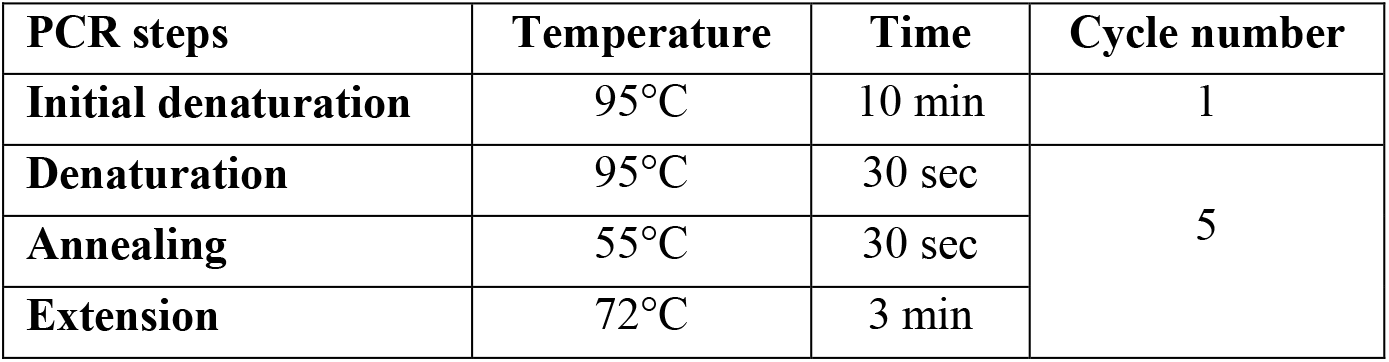
PCR amplification conditions for Zymo Research Quick-16S library preparation (V1-V2 and V3-V4 amplicon sequencing) – PCR II (barcoded PCR)

**DNA from MCS samples**: 2 ng/µl DNA was used to prepare the V1-V2 libraries from microbial mixture

###### Zymo Research V3-V4

The protocol used was the same as described in the ‘*Zymo Research V1-V2*’ section with the following modifications: In the initial PCR step, V3-V4 primers were utilized. **Table 3** provides details of the barcoded primers used, including the pairs and their sequences.

##### PerkinElmer NEXTFLEX® 16S V1-V3 Amplicon-Seq Kit for Illumina

Genomic DNA, having concentrations between 1.6 ng and 36 ng as specified in **Table 5**, was diluted using Nuclease-free Water to maintain a total volume no greater than 36 µL. Subsequently, 12 µL of NEXTflex™ PCR Master Mix and 2 µL of the 16S V1-V3 PCR I Primer Mix were added to the solution. The final reaction volume was adjusted to 50 µL. First amplification step of the PCR cycling was carried out using the settings outlined in **Table 6**.

**Table 5.** Summary table for showing the amount of DNA used for PerkinElmer library preparation, the amount of libraries used for sequencing. The table also shows the barcode ID-s as well as the index sequences.

| Sample | Genomic DNA cc (ng/ul) | Library cc (ng/ul) | Barcode | primer index |
| --- | --- | --- | --- | --- |
| **Q1** | 22,2 | 3,24 | 11 | AAGCGTACGTCC |
| **Q2** | 26,4 | 4,02 | 13 | TCGGGAAGGTCC |
| **Q3** | 25,5 | 8,56 | 37 | TCGGGAAGGTCC |
| **Q4** | 36 | 0,16 | 38 | GAGGCATCGGCC |
| **MN1** | 2,5 | 29,2 | 14 | TTATCAGTCCTT |
| **MN2** | 2,2 | 34,8 | 15 | GTCATCGCGTCC |
| **MN3** | 2,27 | 42,8 | 39 | AATAATTGGTCC |
| **MN4** | 1,6 | 40,6 | 40 | GTCGTCAACCGG |
| **I1** | 3,47 | 34,8 | 16 | CCGTCTCTCCGG |
| **I2** | 4,1 | 5,22 | 18 | ACGCTCTTCCGG |
| **I3** | 2,4 | 41,8 | 41 | AAATCTCAGGCC |
| **I4** | 4,46 | 45,4 | 42 | GTGCGCGGCCGG |
| **Z1** | 4,46 | 25,8 | 13 | AAGCGTACGTCC |
| **Z2** | 3,78 | 13,8 | 14 | TTATCAGTCCTT |
| **Z3** | 3,59 | 34,6 | 16 | CCGTCTCTCCGG |
| **Z4** | 4,42 | 4,1 | 18 | ACGCTCTTCCGG |

**Table 6.** PCR Conditions for PerkinElmer NEXTFLEX® 16S V1-V3 Amplicon-Seq Kit for Illumina – PCR I.

| PCR steps | Temperature | Time | Cycle number |
| --- | --- | --- | --- |
| **Initial denaturation** | 98°C | 4 min | 1 |
| **Denaturation** | 98°C | 30 sec | 8 |
| **Annealing** | 60°C | 30 sec |  |
| **Extension** | 72°C | 30sec |  |
| **Final extension** | 72°C | 4 min | 1 |

PCR cleanup: fifty µL AMPure XP Beads was added to each sample. After mixing, the samples were incubated at room temperature for 5 min. Then, using a magnetic stand, the samples were left until the supernatant clarified. The supernatant was then discarded, and the beads were washed twice with 200 µL of freshly prepared 80% ethanol. Next, the samples were air-dried for 3 min and resuspended in 38 µL of Resuspension Buffer. After a further incubation of 2 minutes at room temperature, 36 µL from the clear supernatant was transferred to fresh tubes. This sample was then subjected to PCR amplification with the addition of 12 µL of NEXTflex™ PCR Master Mix and 2 µL of NEXTflex™ PCR II Barcoded Primer Mix. The procedure was executed following the guidelines specified in **Table 7**. PCR cleanup was carried out in accordance with the purification after the first PCR.

**Table 7.** PCR Conditions for PerkinElmer NEXTFLEX® 16S V1-V3 Amplicon-Seq Kit for Illumina – PCR II.

| PCR steps | Temperature | Time | Cycle number |
| --- | --- | --- | --- |
| **Initial denaturation** | 98°C | 4 min | 1 |
| **Denaturation** | 98°C | 30 sec | varying according to the amount of the initial amount of DNA |
| **Annealing** | 60°C | 30 sec |  |
| **Extension** | 72°C | 30sec |  |
| **Final extension** | 72°C | 4 min | 1 |

###### For the analysis of full-length 16S rRNA gene sequencing

*ONT Rapid Sequencing 16S Barcoding Kit (SQK-RAB204):* Ten ng of high molecular weight genomic DNA (in a 10 ul volume) was used for library preparation from both the canine and MCS samples. DNA isolated using the QIAGEN kit did not meet this criterion. The input DNA was mixed with 14 μl of Nuclease-free water (Invitrogen), 1 μl of 16S Barcode (1 μM; **Table 8**)) and 25 μl of LongAmp Taq 2X master mix (New England Biolabs). PCR Amplification of the samples was carried out according **Table 9**. Amplified DNA samples were transferred to clean 1.5 ml Eppendorf DNA LoBind tubes and mixed with 30 μl of resuspended AMPure XP beads (Beckman Coulter). Next, they were incubated on a Hula mixer for 5 minutes at room temperature. Tubes were placed on a magnetic rack then the supernatant was discarded. The beads were washed with 200 μl of freshly prepared 70% ethanol. Ethanol was removed and the washing was repeated once. After air drying of the beads, samples were removed from the magnet and beads were resuspended in 10 μl of 10 mM Tris-HCl pH 8.0 with 50 mM NaCl. After 2 min incubation at room temperature, samples were placed on the magnet. Ten μl of the clean supernatant, containing the ONT libraries, was transferred to a new Eppendorf DNA LoBind tube. Barcoded libraries were pooled in equal molar ratio and then, 1 μl of RAP was added. The reaction was incubated for 5 minutes at room temperature. One hundred fmoles were loaded on a MinION flow cell.

**Table 8.**
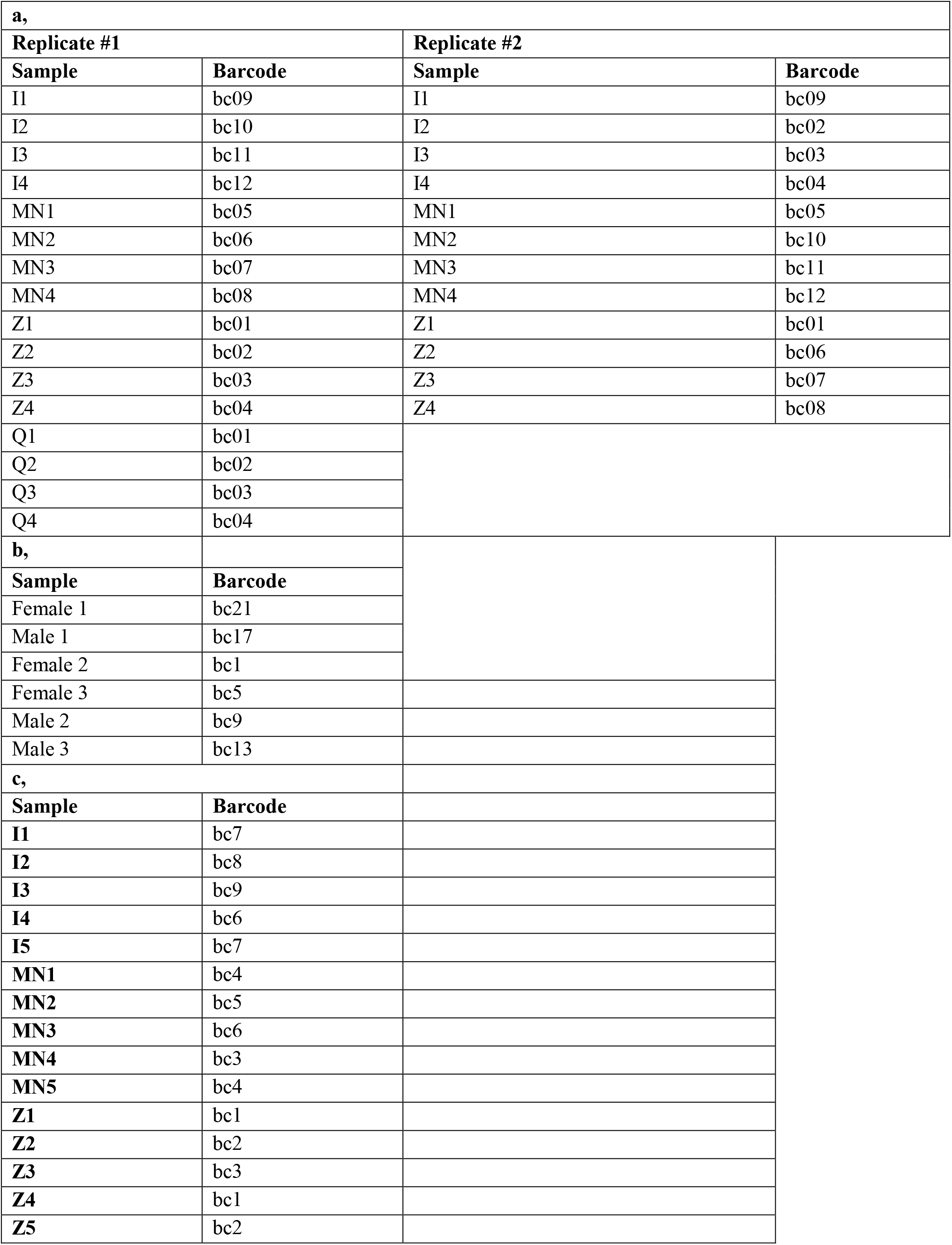
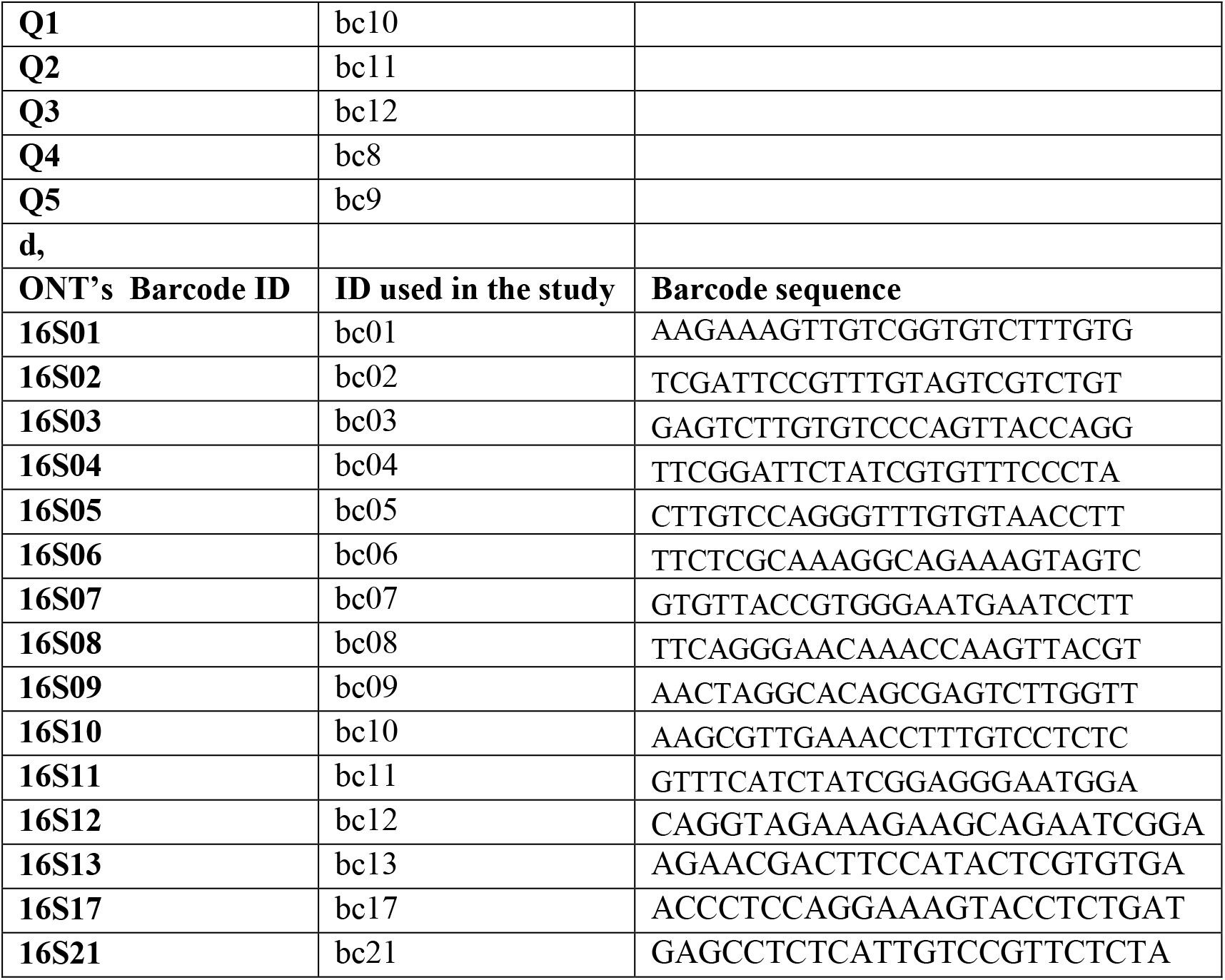
Summary table for showing the barcode IDs and sequences for ONT 16S library preparation. IDs used for library preparation **a,** from the main subject of the study; **b,** from the six additional dogs; **c,** form MCS; **d,** barcode sequences

**Table 9.** PCR Conditions for preparation of ONT V1-V9 libraries.

| PCR step | Temperature | Time | No. of cycles |
| --- | --- | --- | --- |
| **Initial denaturation** | 95 °C | 1 min | 1 |
| **Denaturation** | 95 °C | 20 secs | 25 |
| **Annealing** | 55 °C | 30 secs | 25 |
| **Extension** | 65 °C | 2 mins | 25 |
| **Final extension** | 65 °C | 5 mins | 1 |
| **Hold** | 4 °C | ∞ |  |

*PacBio Full-Length 16S Library Preparation Using SMRTbell Express Template Prep Kit 2.0 Sequel IIe System ICS v10.0 / Sequel II Chemistry 2.0 / SMRT Link v10.0*

For each sample, 1.5 µL of PCR-grade water and 12.5 µL of 2X KAPA HiFi HotStart ReadyMix were mixed. Subsequently, 3 µL of barcoded forward primer solution (2.5 µM, see sequences in **Table 10**) was added. This was followed by the addition of 3 µL of the respective reverse primer solution (**Table 10**) and 5 µL of the DNA sample. DNA amplification was carried out according to the parameters listed in **Table 11**.

**Table 10.**
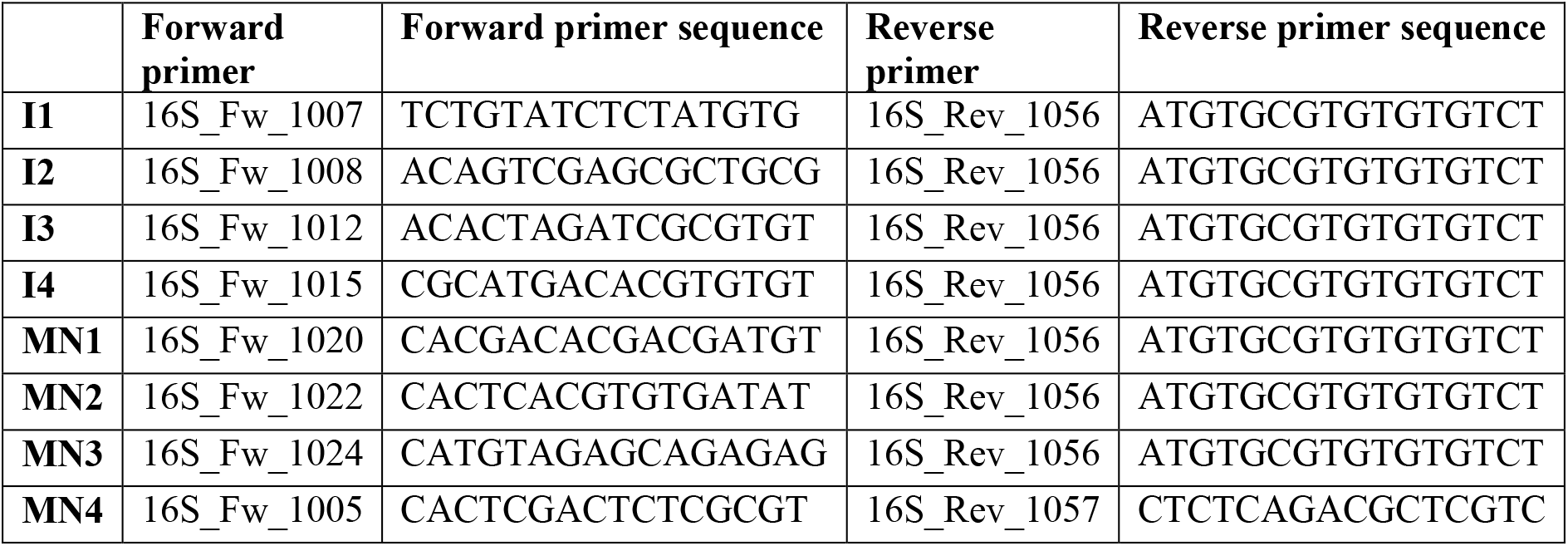
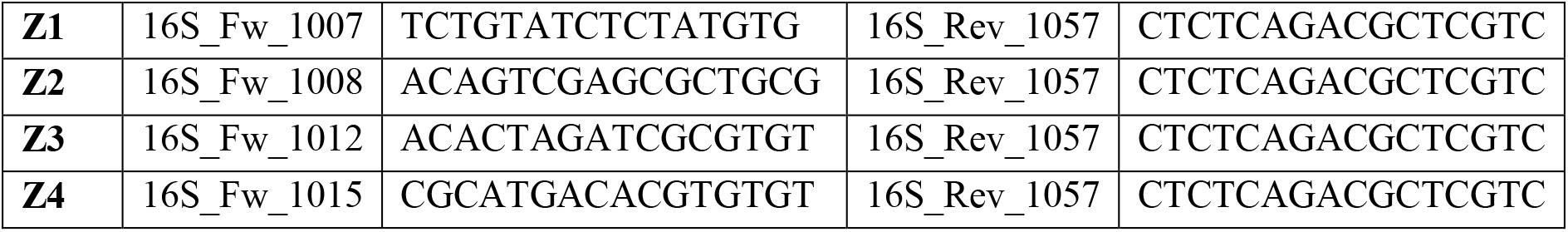
Summary table of primers used for PacBio 16S library preparation.

**Table 11.** PCR conditions for preparation of PacBio V1-V9 libraries–.

| PCR steps | Temperature | Time | Cycle number |
| --- | --- | --- | --- |
| **Initial denaturation** | 95°C | 3 min | 1 |
| **Denaturation** | 95°C | 30 sec | 25 |
| **Annealing** | 57°C | 30 sec |  |
| **Extension** | 72°C | 60sec |  |

###### Illumina DNA Prep

Sixty ng of DNA (in 30 µl) was used as total input per sample. Ten µl Tagmentation Buffer 1 (TB1) was mixed with 10 µl Bead-Linked Transposomes (BLT), and 20 µl of this mixture was added to a DNA sample. The mixture was incubated at 55°C for 15 min and then held at 10 °C. Following this, 10 µl of Tagment Stop Buffer (TSB) was added to the sample and gently mixed.

The samples were incubated at 37°C for 15 min, and then kept at 10°C. Next, the samples were placed on a magnetic stand for 3 min, and the supernatant was discarded. The sample was removed from the magnet and 100 µl of Tagment Wash Buffer (TWB) was slowly added directly onto the beads. The sample was placed back on the magnetic stand, the supernatant was discarded, and the wash step was performed again. A mixture of 20 µl of Enhanced PCR Mix (EPM) and 20 µl of nuclease-free water was prepared, and 40 µl of this mixture was added to the washed beads. Index adapters (i5 and i7, 5 µl each; see **Table 12**) were added, and PCR was carried out according to **Table 13**. After amplification, the libraries were cleaned up. First, the samples were placed on a magnet for approximately 5 min. Forty-five µl of supernatant from each PCR product was transferred to a new tube. Forty µl of nuclease-free water and 45 µl of Sample Purification Beads (SPB) were added to the supernatant and the samples were mixed at 1600 rpm for 1 min. They were incubated at room temperature for 5 min. After this step, the samples were placed on a magnet, and 15 µl of SPB were added to new tubes. Next, 125 µl of supernatant from each sample was added to the tubes containing 15 µl of undiluted SPB. The samples were mixed at 1600 rpm for 1 min, and then incubated at room temperature for 5 min. The supernatant was discarded, and the washing step was carried out twice with 200 µl of freshly prepared 80% ethanol. The sample was stored on magnetic stand for 30 secs, and then the ethanol was removed. After the second washing step, the pellet was air dried. Next, 32 µl RSB was added to the beads, and they were mixed and incubated for 2 min. Finally, the sample was placed on a magnetic stand for 2 min, and the supernatant containing the prepared library was transferred to a new tube.

**Table 12.**
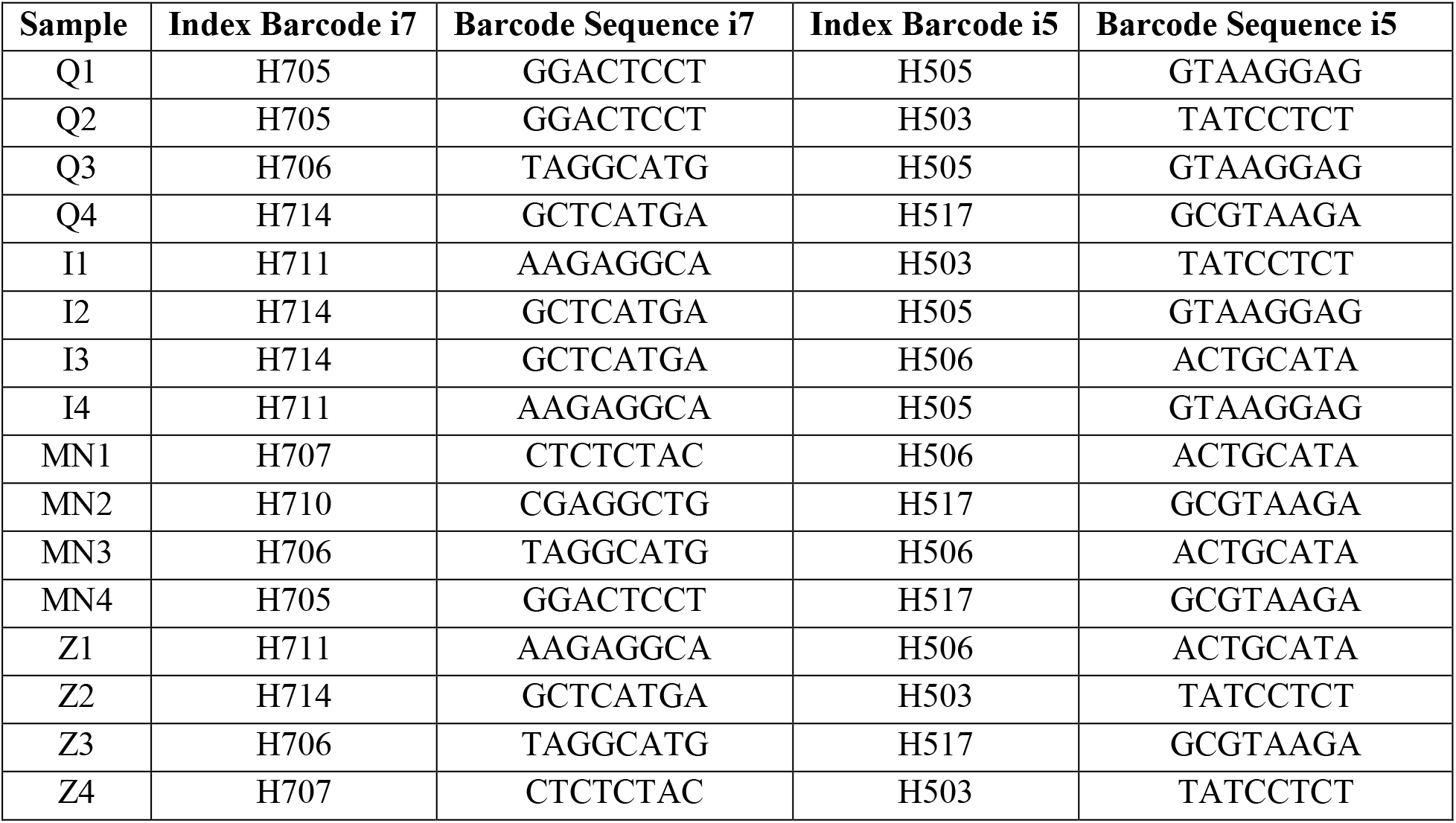
Summary table of primers from Illumina DNA Prep Kit used for WGS library preparation.

| Sample | Index Barcode i7 | Barcode Sequence i7 | Index Barcode i5 | Barcode Sequence i5 |
| --- | --- | --- | --- | --- |
| Q1 | H705 | GGACTCCT | H505 | GTAAGGAG |
| Q2 | H705 | GGACTCCT | H503 | TATCCTCT |
| Q3 | H706 | TAGGCATG | H505 | GTAAGGAG |
| Q4 | H714 | GCTCATGA | H517 | GCGTAAGA |
| I1 | H711 | AAGAGGCA | H503 | TATCCTCT |
| I2 | H714 | GCTCATGA | H505 | GTAAGGAG |
| I3 | H714 | GCTCATGA | H506 | ACTGCATA |
| I4 | H711 | AAGAGGCA | H505 | GTAAGGAG |
| MN1 | H707 | CTCTCTAC | H506 | ACTGCATA |
| MN2 | H710 | CGAGGCTG | H517 | GCGTAAGA |
| MN3 | H706 | TAGGCATG | H506 | ACTGCATA |
| MN4 | H705 | GGACTCCT | H517 | GCGTAAGA |
| Z1 | H711 | AAGAGGCA | H506 | ACTGCATA |
| Z2 | H714 | GCTCATGA | H503 | TATCCTCT |
| Z3 | H706 | TAGGCATG | H517 | GCGTAAGA |
| Z4 | H707 | CTCTCTAC | H503 | TATCCTCT |

**Table 13.** PCR Conditions for preparation of Illumina WGS libraries.

| PCR steps | Temperature | Time | Cycle number |
| --- | --- | --- | --- |
| **Pre-heating** | 68°C | 3 min | 1 |
| **Initial denaturation** | 98°C | 3 min | 1 |
| **Denaturation** | 98°C | 45 sec | 5 |
| **Annealing** | 62°C | 30 sec |  |
| **Extension** | 68°C | 2 min |  |
| **Final extension** | 68°C | 1 min | 1 |

### SEQUENCING

Altogether nine MiSeq Reagent Kit v2 and four MiSeq Reagent Kit v2 Nano were used for SRS (**Table 14**). V1-V9 sequencing was carried out on ONT MinION and on PacBio Sequel IIe instruments. ONT V1-V9 barcoded libraries from the canine samples were loaded onto three MinION flow cells and two additional flow cells were run for sequencing the MCS samples. PacBio V1-V9 libraries were barcoded and run on a Sequel SMRT Cell 8M.

**Table 14.**
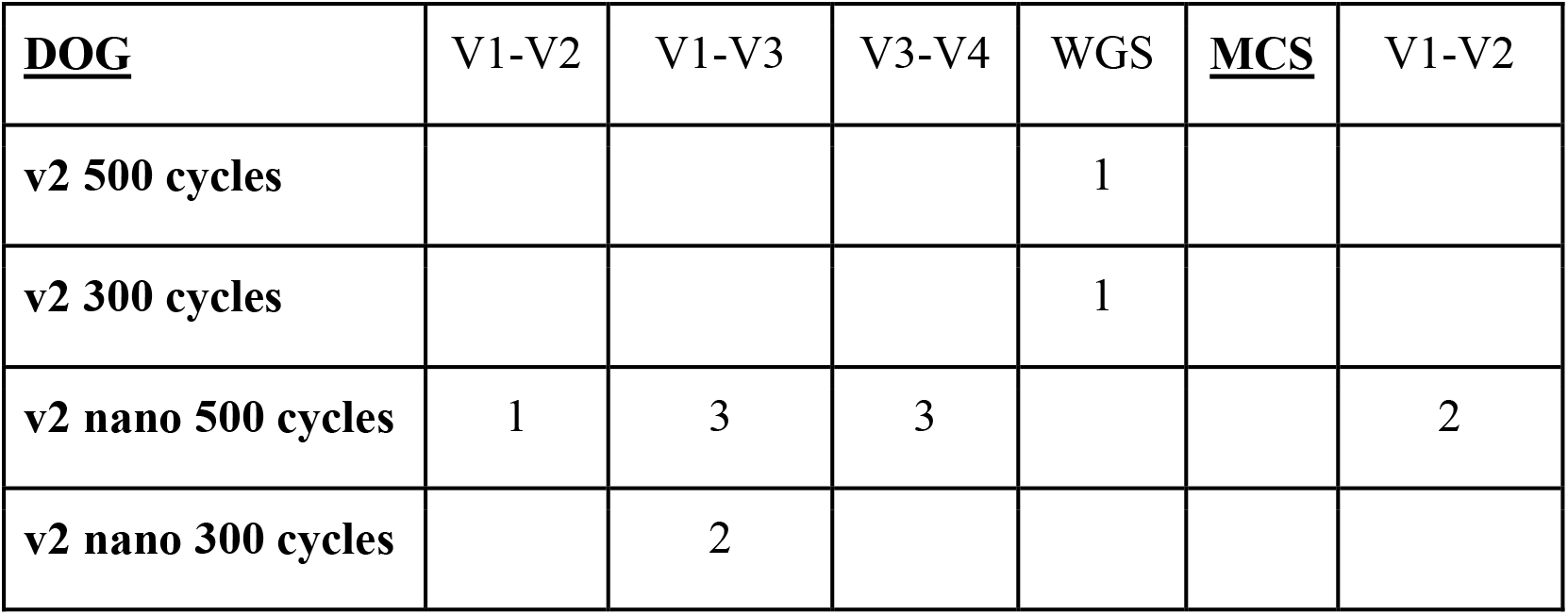
Illumina Reagent Kits used for this study.

| **<u>DOG</u>** | V1-V2 | V1-V3 | V3-V4 | WGS | **<u>MCS</u>** | V1-V2 |
| --- | --- | --- | --- | --- | --- | --- |
| **v2 500 cycles** |  |  |  | 1 |  |  |
| **v2 300 cycles** |  |  |  | 1 |  |  |
| **v2 nano 500 cycles** | 1 | 3 | 3 |  |  | 2 |
| **v2 nano 300 cycles** |  | 2 |  |  |  |  |

#### BIOINFORMATIC ANALYSIS

##### Development of a bioinformatic tool

We developed minitax a versatile taxonomic assignment tool designed to address the challenges posed by a diverse range of sequencing data types. This program uses minimap2, chosen for its speed and accuracy, to align the sequencing reads to the specified reference database. The minitax uses minimap2 platform-specific parameter settings for the initial read alignment. It then loads the alignments into an R environment using Rsamtools^64^ and merges the alignments with the database. Subsequently, it performs sophisticated post-alignment processing steps to find the best alignment for each read based on mapping quality (mapq) and CIGAR scores. These CIGAR scoring schemes and other parameters are controllable by the user. Subsequently, it summarizes the read counts for each taxonomic level. Finally, it creates a report from the summary statistics and plots the results.

#### Analysis of the raw data

##### Illumina V1-V2, V3-V and V1-V3

*DADA2 pipeline* on Illumina amplicon reads: The raw reads were subjected to DADA2^45^ (version 1.22) for quality control, filtering, trimming and subsequent dereplication (ASV generation), bimera detection/removal and lastly the taxonomic assignment. The latter was carried out using either the SILVA 16S database^65^ (version: 138.1) or Emu’s database (version: v3.4.4). The exact parameters and settings can be found in SUPPTAB_X (dada2_config.tsv) and the complete workflow on the GitHub page (dada2.WF.R)

*Emu* and *minitax pipeline* on Illumina reads: The Emu software^47^ (version: v3.4.4) and the in-house developed minitax program (https://github.com/Balays/minitax) (version 1.0) were used on the raw Illumina reads with the following parameters: *--type sr* and *--N 10,* otherwise default settings was applied. We used Emu’s default database in both cases and additionally a genome collection from NCBI in the case of minitax. Both programs utilize minimap2^49^ for initial read alignment.

##### ONT V1-V9

The raw voltage values obtained from the MinION sequencer platform were basecalled with the 6.1.5 version of Guppy (MinKNOW 20.05.8) at super-high accuracy. The reads were demultiplexed using the SQK-RAB204 barcode sequences. During basecalling, the minimum quality score was set to 8, thus fail and pass reads were obtained. For further analyses only the pass reads were used. The pass reads were then applied as an input for Emu (version: v3.4.4) and minitax (version 1.0) using the following parameters: *--type map-ont* and *--N 10* otherwise defaults, and using Emu’s default database for both programs and additionally a genome collection from NCBI in the case of minitax. In addition, ONT’s EPI2ME pipeline (3.6.1) was also used

##### PacBio V1-V9

Basecalling, demultiplexing and generation of HiFi reads were carried out by using the SMRT Link 10.2.0.133434 analysis software from PacBio. High quality CCS reads were filtered: --min- qv 20. The filtered reads were then used as an input for Emu (version: v3.4.4) and minitax (version 1.0) using the following parameters: --type map-pb and --N 10 otherwise defaults, and using Emu’s default database in both cases and additionally a genome collection from NCBI in the case of minitax.

##### Illumina WGS

The raw reads were trimmed using Trim Galore and host reads filtered with (https://github.com/FelixKrueger/TrimGalore) BMTagger (https://hpc.ilri.cgiar.org/bmtagger-software) using the *Canis lupus familiaris* genome assembly (GCF_014441545.1). The quality controlled reads were subjected to either the fast and sensitive taxonomic classifier program sourmash (version 4.8.2) using the provided Genbank genomes database (from March 2022) or the minitax software (version: 1.0) using default parameters and a genome collection from NCBI as a reference.

##### Benchmarking Datasets

Besides the Zymo D6300 Microbial Community Mixture sequenced using ONT V1-V9 and Illumina V1-V2 methods described earlier, we evaluated the performance of minitax on two other datasets published elsewhere.

PacBio HiFi WGS of Zymo MCM D6331 Microbial Community: This dataset was used in Portik et al. 2022, for benchmarking of long-read mWGS software. The *fastq* files were downloaded from NCBI (NCBI accession: SRX9569057) and used as an input for either the sourmash program or minitax.

CAMISIM Datasets: The CAMISIM Simulated Mouse Gut Project^44^, provided simulated reads in fastq format, of which we randomly downloaded 10 samples each for PacBio and Illumina to use as input for the programs.

Performance Metrics: To provide a deep insight into the capabilities of minitax relative to references, precision, recall, F1 and F0.5 scores were calculated against the theoretical compositions for 1%, 0.1%, and 0.01% species detection thresholds after *Portik et al., 2020*^33^. Additionally, we calculated Pearson correlations between the theoretical and observed compositions and the associated r^2^ values for each taxonomic level.

##### Databases

We utilized a comprehensive set of ∼18,000 genomes spanning 13,000 species of *Bacteria*, *Archaea* and *Eukaryotes*, downloaded in February, 2022 from NCBI as a reference for both WGS and 16-Seq data. Additionally, for the amplicon sequencing datasets, we employed either the Emu or the SILVA (version 138) 16S databases, depending on the software used.

##### CAMISIM

The simulated mouse gut dataset was used for additional validation of the minitax workflow. Fifteen samples for both the short-read (Illumina) and long-read (PacBio) datasets were employed.

##### Downstream data analysis

The results of the different programs were combined into ‘phyloseq’ objects^66^ and subsequently combined into a single, comprehensive phyloseq object encompassing all the metagenomic read count data. Subsequent data analysis was performed using this phyloseq object in the R environment, with the addition of these packages: tidyverse^67^, FactoMineR^68^, and vegan^69^. The scripts for the complete downstream data analysis and plots can be downloaded from GitHub repository (https://github.com/Balays/Microbiome-Method-Comparison).

##### Downsampling

The Kaiju software package was run on the Illumina mWGS data, and using the output obtained, the number of reads in the sample was randomly reduced to the following values: 2500000, 2000000, 1500000, 1000000, 750000, 500000, 250000, 225000, 200000, 175000, 150000, 125000, 100000, 75000, 50000, 20000, 10000 and 5000. The resulting data were assigned a taxonomic classification. The data were then analyzed with the phyloseq R software package and visualized with ggplot2.

The raw reads from the non-WGS dataset were subsampled using SeqTK, randomly selecting subsets of 2,500,000; 2,000,000; 1,500,000; 1,000,000; 750,000; 500,000; 250,000; 225,000; 200,000; 175,000; 150,000; 125,000; 100,000; 75,000; 50,000; 20,000; 10,000; and 5,000 reads. The subsampled datasets were then processed using EMU with default settings. Subsequent taxonomic composition analysis was performed using Phyloseq, and the results were visualized using ggplot2.”

## Data availability

European Nucleotide Archive: PRJEB59610

## Code availability

minitax: https://github.com/Balays/minitax

other in-house scripts

- https://github.com/Balays/Microbiome-Method-Comparison
- https://github.com/gabor-gulyas/Technical-article-downsample

DADA2: https://benjjneb.github.io/dada2/

Emu: https://gitlab.com/treangenlab/emu

Trim Galore: https://github.com/FelixKrueger/TrimGalore

BMTagger: https://hpc.ilri.cgiar.org/bmtagger-software

## ACKNOWLEDGEMENTS

This work was supported by the Momentum program I of the Hungarian Academy of Sciences LP2020-8/2020 (D.T.), the National Research, Development and Innovation Office FK 142676 (D.T.) and K 142674 (Z.B.). Á.D. was a grantee of the New National Excellence Program (ÚNKP-22-3-SZTE-245).

## AUTHOR CONTRIBUTIONS

Z. B., B. K. and D.T. conceived the project and designed experiments. G. G., Á. D., T. J., Z. C. and D. T. performed experiments. G. G., K. B., I. P., Z. B. and D. T. conducted computational analyses. D. T. supervised the project. Z.B. and D.T. contributed key reagents. B. K., M. M. H., Z. B. and D. T. wrote the manuscript. All authors participated in the review and editing of the manuscript.

## COMPETING INTERESTS

The authors declare no competing interests

## LEGEND TO SUPPLEMENTARY FIGURES

**Supplementary Fig. 1: Comparative analysis of cost and hands-on time for DNA isolation kits.**

This figure contrasts the cost per sample with the hands-on time required for different DNA isolation kits, with prices presented in USD and based on the summer 2023 sales in Hungary.

**Supplementary Fig. 2: Analysis of canine stool DNA extraction using Qiagen kit and Agilent ScreenTape Assay.**

The figure illustrates DNA extracted from control canine fecal samples utilizing the Qiagen kit, which is then analyzed using the Agilent genomic ScreenTape assay. Lane A1 (L) represents the DNA ladder. Abbreviations are as follows: y: years; w: weeks.

**Supplementary Fig. 3: Correlation between downsampling and diversity.**

The relationship between read count and microbial diversity is explored in this figure, underscoring the opportunity for cost-efficiency in shotgun sequencing. Even with a reduction to approximately 200,000 reads, the diversity across samples remains stable. For assessments focusing on bacterial composition rather than detection of rare species, a read count of 200,000 per sample is sufficient.

**Supplementary Fig. 4: β-diversity assessment using Bray-Curtis distance and PERMIDISP analysis**

**a,** The PERMIDISP results display a multivariate analysis of group dispersion homogeneity, indicating the distance to the centroid for each sample in relation to DNA isolation kits and library preparation protocols.

**b,** The PCoA results depict the variance and resemblance among the DNA isolation protocols for each library preparation.

**c,** The results of the PERMIDISP analysis are presented, categorized by library preparation protocols and DNA isolation kits.

**Supplementary Fig. 5: TapeStation analysis of various DNA libraries from canine stool samples.**

This figure contrasts the cost per sample and the hands-on time associated with various library preparation kits, denoted in USD, based on the summer 2023 sales prices in Hungary.

**Supplementary Fig. 6: The TapeStation images of the various libraries. a-j, represent samples from the primary test dog;**

**k-l,** indicate samples from the control dogs.

**a-c ,** 16S V1-V3 libraries created using the PerkinElmer NEXTFLEX® Kit and analyzed with the Agilent D1000 ScreenTape assay. Lanes EL1 (L) represent electronic DNA ladders. R indicates a technical replicate (Dog stool).

**d,** Zymo Research V1-V2 libraries examined with the Agilent D1000 ScreenTape assay. Lane E1 (L) represents the electronic DNA ladder.

**e-g,** Shotgun (mWGS, Illumina DNA Prep) libraries derived from canine stool samples.

**h-j,** V1-V9 library pools from dog stool prepared for ONT (h) and PacBio (i) sequencing. Figure h displays a pool of sixteen libraries, with four each prepared from DNA isolated using Q, I, MN, and Z DNA purification kits for ONT sequencing. Figure i shows twelve PacBio libraries (four each from I, MN, and Z-prepped DNAs). The PCR products were approximately 1.5 kb in size. Abbreviations: MN: Macherey-Nagel, I: Invitrogen, Z: Zymo Research.

**k-l,** V1-V3 libraries made using the PerkinElmer NEXTFLEX® 16S Amplicon-Seq Kit and analyzed with the Agilent D1000 ScreenTape assay. Lane E1 (L) is the electronic DNA ladder, and A1 (L) is the ladder.

**Supplementary Fig. 7: Comparison of the bacterial composition of various samples at the phylum level.**

This figure presents a comparison of bacterial composition at the Phylum level in various samples. **a,** Tables show the proportions of the most abundant Phyla in samples obtained using different DNA extraction and library preparation methods.

**b,** Bar charts highlight the discrepancies between the proportions of the most abundant bacterial Phyla from other groups’ data and our datasets.

**Supplementary Fig. 8: Ratio of Gram-Positive and Gram-Negative Bacteria in Canine Stool as a Result of Various Laboratory Methods**

The barplots show the ratio of Gram(+) and Gram(-) bacteria in each sample, according to library preparation protocols and DNA isolation methods. Zymo’s results indicate a slight overrepresentation of Gram negatives compared to the Macherey-Nagel and Invitrogen methods, with the Qiagen method showing a significant overrepresentation.

**Supplementary Fig. 9: Agilent ScreenTape Assay: comparing DNA yields and fragment lengths from MCS samples**

The images depict the Agilent genomic DNA ScreenTape assay results for DNA extracted from MCS. Lanes A1 (L) represent DNA ladders. The Invitrogen and Qiagen samples showed lower DNA amounts compared to others, with Zymo yielding the highest amount. Despite Zymo Kit being designed for HMW DNA isolation, Invitrogen and Qiagen yielded longer DNA fragments from Zymo MCS samples.

**a,** First three replicates.

**b,** Two subsequent replicates.

**Supplementary Fig. 10: Evaluating minitax: a comparison with other methods based on Pearson’s correlations.**

This figure compares minitax with other methods based on the Pearson’s correlations between theoretical and observed compositions (r2 values).

**a,** Comparison with Emu on ONT V1-V9 Sequencing of Zymo D3600 MCS data.

**b,** Comparison with DADA2 and Emu on Illumina V1-V2 Sequencing of Zymo D3600 MCS.

**c,** Comparison with sourmash on PacBio HiFi WGS of Zymo MCS D6331.

## Notes

### Competing Interest Statement

The authors have declared no competing interest.

